# Essential role of non-vesicular lipid transport in microtubule-controlled cell polarisation

**DOI:** 10.1101/2020.11.09.374702

**Authors:** M. Hersberger-Trost, D. Dreher, S.M. Huisman, A.R. Kijowski, M. Gemünden, R.W. Klemm, D. Brunner

**Affiliations:** Department of Molecular Life Sciences, University of Zurich, Winterthurerstrasse 190, 8057 Zurich, Switzerland

## Abstract

Cell polarisation is a fundamental biological process. Fission yeast is a key model system to study the molecular basis of microtubule-controlled cell polarisation. In this process, cells define prospective growth sites by generating distinct plasma membrane domains enriched in *de novo* synthesised sterols. Microtubules restrict the number and location of these domains by depositing factors at the cell poles. The mechanisms underlying such sterol-rich membrane domain formation and polarisation are largely unknown. We found that the oxysterol-binding proteins kes1p, osh2p and kes3p define three independent sterol delivery pathways to the plasma membrane. These mediate different phases of cell polarisation in a phosphoinositide-dependent fashion and differ in their requirement for vesicular trafficking steps. The redundant, kes1p- and osh2p-dependent pathways are vital and prime cell polarisation by mediating the formation of randomly distributed sterol-rich plasma membrane domains. Subsequent microtubule-controlled polarisation of these domains preferentially employs kes1p that directly delivers sterols to the plasma membrane independent of cdc42p. In cells lacking kes1p, polarisation becomes cdc42p-dependent, utilising mainly the kes3p-dependent pathway. Our study uncovers an essential biological function for non-vesicular lipid transport and establishes a molecular basis for different sterol-delivery pathways acting in cdc42p-independent and cdc42p-dependent cell polarisation.

## Introduction

Cell polarisation, the act of symmetry breaking, is a fundamental process in all cells. It can be triggered cell autonomously by inherent molecular systems, or by externally stimulated signalling pathways. The common response is a cell type specific re-arrangement of the cytoskeleton and the asymmetric organisation of a number of cellular components. In many cell types, polarity was shown to depend on the activity of the GTPase Cdc42 ^1–5^. In budding yeast, constitutively activated Cdc42 spontaneously imposes symmetry breaking in cells lacking the usual polarity signals, leading to the interpretation that polarity signals act by directly controlling the positioning of Cdc42 activating guanine nucleotide exchange factors (GEFs) ^6–10^.

In various cell types, microtubules were found to provide a signal that mediates proper cell polarisation ^11–14^ The underlying molecular mechanisms are not well understood but it is generally assumed that this cell intrinsic pathway directly positions the GEFs controlling Cdc42 activation ^1,3^. This is also the case for growth polarisation in fission yeast ^15^. During proliferation, these cylindrical cells position growth zones at their poles via their interphase microtubules in combination with a microtubule-independent polarity inheritance mechanism that so far has not been molecularly described ^16^.

Polarity inheritance plays no role in cells polarising *de novo* when exiting a starvation period ^17^ Instead, such cells now fully rely on microtubules and the polarity factor tea1p. In the absence of a functioning microtubule/tea1p system such cells show completely randomised growth site positioning. Starved, quiescent cells thus provide a powerful experimental system to dissect the molecular mechanisms underlying microtubule-mediated control of cell polarisation. Microtubule/tea1p-controlled cell polarisation was found to follow a stereotypical, linear sequence of events that can be broken down into four phases (P1-P4) (Fig. 1A). The process fully depends on the *de novo* biosynthesis of sterols. In P1, these sterols initially enrich in multiple, randomly distributed, sterol-rich membrane (SRM) domains at the plasma membrane. Each domain recruits established polarity- and growth factors and can in principle develop into a site of growth. SRM domain formation occurs independent of the microtubules that simultaneously self-organise in a bipolar fashion, aligned with the long cell axis ^18^. This results in the deposition of the microtubule plus end-tracking protein tea1p at both cell poles ^19^. The actual polarisation occurs in P2 ^17^ where the polar position of tea1p ensures the expansion and maintenance of SRM domains at cell poles. At the same time all SRM domains in the cell middle disappear ^17^, confining subsequent cell growth initiation to the poles. Interestingly, this key polarisation step was found to occur independent of Cdc42 (termed cdc42p in fission yeast). Cdc42p activity only becomes detectable towards the end of P2 ^17^ In P3, one of the two predetermined poles activates growth, and in P4 cells start growing at the other pole as well. Unlike polarisation in P2, polarised growth in P3 is fully cdc42p dependent. Thus, *de novo* cell polarisation in this organism, employs cdc42p to maintain predefined cell polarity.

**Fig. 1:**
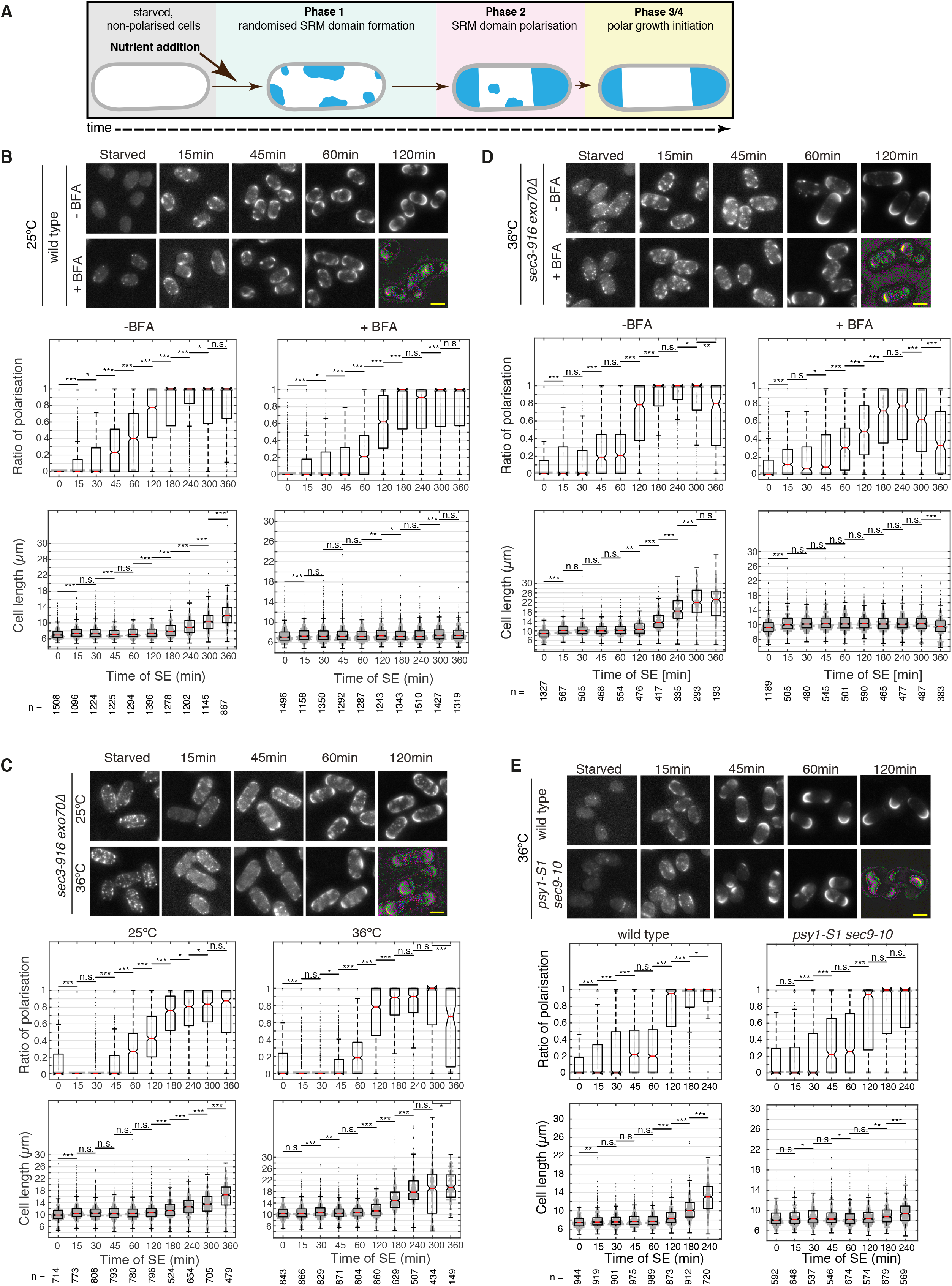
Sterol delivery occurs independent of vesicular transport. (**A**) Cartoon depicting the 4 phases of SE in fission yeast ^17^ (**B)** SRM domain formation and polarisation during SE at 36°C of filipin-stained wild type cells treated with BFA dissolved in DMSO (+BFA) and DMSO alone (- BFA control). Notched box plots in this and all following figures show the polarisation rates at different time points during SE. The value distribution is shown in grey. Asterisks represent p-values to help assessing the changes in SRM domain dynamics and growth initiation that correlate to the progression of cells through the phase of SE, from timepoint to timepoint. * represents p≤0.05, ** represents p≤0.01, *** represents p≤0.001 and “n.s.” stands for non-significant difference (Supplementary Materials). (**C**) SRM domain formation and polarisation during SE at permissive (25°C) and restrictive temperatures (36°C) of filipin-stained *sec3-916 exo70*Δ double mutant cells. Notched box plots show the polarisation rates and cell lengths at different time points during SE with p-values. (**D**) SRM domain formation and polarisation during SE at restrictive temperatures (36°C) of filipin-stained *sec3-916 exo70*Δ double mutant cells treated with DMSO (- BFA control) and BFA in DMSO (+BFA). Notched box plots show the polarisation rates and cell lengths at different time points during SE with p-values. (**E**) SRM domain formation and polarisation during SE at restrictive temperatures (36°C), comparing filipin-stained wild type and *psy1-S1 sec9-10* double mutant cells. Notched box plots show the polarisation rates and cell lengths at different time points during SE with p-values. Scale bars: 5μm.

*Teal* deleted cells (*teal*Δ) enter P1 normally but unlike wild type cells, they prolong this state for a variable period of time ^17^ The cells then skip P2 and proceed straight into P3, producing a single growth site originating from any of the randomly positioned SRM domains formed in P1. Depending on the location of the chosen domain, cells will grow branched or bent, and occasionally even straight. Growth initiation in *tea1*Δ cells coincides with the removal of all other SRM domains. Consequently, cells no longer possess a second, prospective growth site, explaining why the mutants never enter P4. In the wild type, in contrast, polar positioned tea1p maintains the second, future growth site throughout mono-polar growth in P3.

These findings raised the question, how SRM domains form in P1, and how tea1p confines their polar localisation in P2. Here we used functional perturbation experiments to explore the molecular basis of sterol cell surface delivery for the initial SRM domain formation during P1, and for microtubule/tea1p-mediated SRM domain polarisation during P2. Our results show that in P1, SRM domain formation occurs by two alternative delivery routes. The first involves the sterol transporter kes1p that delivers sterols to the plasma membrane independent of vesicular transport carriers. The second requires another non-vesicular sterol transporter, osh2p, but also the two t-SNAREs mediating vesicular fusion at the plasma membrane. Together, these routes are absolutely essential for the survival of cells exiting starvation although they are fully dispensable in proliferating cells.

When SRM domains polarise in P2, cells preferentially use the kes1p sterol delivery route. In the absence of kes1p, cells still polarise in a tea1p-controlled manner, but now via cdc42p and a third sterol delivery route, involving the sterol transporter kes3p. Altogether, our results show that in *de novo* polarising cells, sterol delivery to the plasma membrane can occur via three different routes, which are differentially deployed during P1 and P2 of cell polarisation. Our findings not only disentangle the role of these pathways in the critical steps of cell polarisation, but also clarify their contribution to cdc42p-independent and cdc42p-dependent polarity mechanisms.

## Results

### SRM domain polarisation occurs independent of vesicular trafficking

Sterols are produced in the endoplasmic reticulum (ER). An established route for sterol delivery from the ER to the plasma membrane is through classical vesicular transport via the Golgi apparatus. Post Golgi secretory vesicles then fuse with the plasma membrane in a SNARE-dependent manner ^20–22^. In polarising fission yeast, this would be an obvious route for cell surface delivery of *de novo* synthesised sterols required for SRM domain formation in P1. In P2, the microtubule/tea1p system would then mediate SRM domain polarisation by restricting the delivery to the cell poles. To test this assumption, we carried out functional perturbation experiments using small molecule inhibitors and genetics on polarising cells during starvation exit (SE). We developed a simple automated image analysis protocol for unbiased quantification of the degree of cell polarisation in wild type cells and various mutants (Methods). To first test whether vesicular sterol delivery requires trafficking via the Golgi apparatus we treated cells with Brefeldin A (BFA), a drug that interferes with Golgi integrity (Methods) ^23^. We confirmed immediate drug action in cells exiting starvation, by showing that BFA treatment efficiently blocked the delivery of the multi-pass transmembrane protein tna1p tagged with GFP (Fig. S1A). As previously shown, GFP-tna1p accurately labels SRM domains in the PM during all phases of SE ^17^ In the presence of BFA very little GFP-tna1p reached the cell membrane and instead the protein started to accumulate intracellularly with a distribution that is typical for markers of the ER (Fig. S1A) ^24^ Using the sterol-binding dye filipin, we found that BFA neither affected SRM domain formation in P1 nor SRM domain polarisation in P2, suggesting that these phases do not depend on Golgi function (Fig. 1B) ^25^. In P3, BFA treated cells were unable to initiate growth, consistent with effective disruption of Golgi function (Fig. 1B). To further investigate for a function of vesicular sterol delivery, we tested whether *de novo* cell polarisation is defective in strains carrying temperature-sensitive alleles of *sec3, sec8, ypt2* and *pob1*, encoding essential components of exocytosis ^26–30^. At 36°C, the mutant’s restrictive temperature, the fluorescence of our secretory protein marker GFP-tna1p was strongly impaired precluding its use as a live marker of SRM domains in ts-mutant analysis (Fig. S1C). Instead, we analysed cell polarisation by monitoring the localisation of filipin-stained SRM domains. We have previously shown that in standard growth conditions filipin is a reliable marker of SRM domain position and cell polarisation under our strict experimental conditions ^17^. To confirm that this is true also for cells exiting starvation at 36°C we stained wild type cells taken from a culture at defined, consecutive time points during SE. Cells exited starvation 36°C following the same sequence of events and with similar timing as at 25°C. This shows that filipin staining reliably monitors SRM domain formation and cell polarisation also at these higher temperatures (Fig S1B). To observe SE of temperature sensitive mutants under fully mutant conditions, we shifted starved cells to 36°C one hour before initiating SE, and kept them at this temperature throughout the experiment. Despite the strong block in secretion in the aforementioned mutants, P1 and P2 still occurred with kinetics similar to wild type control cells (Fig. S1D). This is consistent with previous results suggesting that, to a limited extent, cells harbouring these mutant alleles can continue polar growth, while the most sensitive phenotype is the completion of cytokinesis ^30,31^09/11/2020 11:36:00. A possible explanation for this observation is functional redundancy amongst exocyst components, as shown e.g. for exo70p and sec3p in the late secretory pathway ^28,31^. Consistently, haploid spores that were mutant for both proteins failed to germinate ^31^. To test such effective interference with exocyst function during SE, we generated cells carrying the *exo70* deletion and the *sec3-916* temperature sensitive mutant (*exo70*Δ *sec3-916*). Interestingly, *exo70*Δ *sec3-916* cells that were glucose starved at 25°C, the permissive temperature, displayed prominent filipin-positive punctae all over their plasma membrane prior to SE (Fig. 1C). During SE under permissive conditions, these dots remained visible throughout the polarisation process. However, polarisation and growth initiation of such cells occurred similar to the wild type, except that removal of central SRM domains was somewhat delayed (Fig. 1C). Under such conditions, *exo70*Δ *sec3-916* cells went through the phases of SE very similar to *exo70*Δ *sec3-916* control cells at the permissive temperature of 25°C (Fig. 1C). This suggests that exo70p and sec3p, and thus the late secretory pathway, are dispensable for microtubule-controlled *de novo* cell polarisation. The only obvious difference between *exo70*Δ *sec3-916* cells at 25°C and 36°C occurred at late time points when cells exiting starvation at 36°C started dying. Dying cells displayed strong fluorescence all over the cell, which was interpreted by our analysis tool as depolarised cells and resulted in a drop of the polarisation ratio at 360minutes of SE.

Interestingly, *exo70*Δ *sec3-916* cells exiting starvation at 36°C entered P3 and initiated growth similar to wild type cells (Fig. 1B and C). Only when additionally treating the cells with BFA, did we completely block P3 (Fig. 1D). Importantly, BFA-treatment of *exo70*Δ *sec3-916* cells, did not affect P1 and it only added a slight delay to SRM domain polarisation in P2 (Fig. 1D). This delay is likely due to the lack of growth, as the polarisation ratio peaked with the onset of P3 in the control cells, while it remained lower in BFA treated cells. BFA also enhanced cell death. These results show that the known vesicular trafficking routes are not required to proceed through P1 and P2.

Since our findings cannot exclude the activity of yet another, unknown vesicular transport pathway, we next blocked the last step of exocytic vesicle fusion with the plasma membrane. We generated cells carrying temperature-sensitive mutant alleles of both essential plasma membrane-resident t-SNAREs psy1p and sec9p (*psy1-S1 sec9-10*) ^32^. After having entered starvation at the permissive temperature of 25°C, the filipin staining of *psy1-S1 sec9-10* cells revealed the presence of 8.4% (n=1377) septated cells suggesting that the last cytokinesis before entering quiescence was not completed (Fig. 1E). Nevertheless, all other cells were depolarised similar to starved wild type cells showing that the majority of cells had entered normal quiescence. When *psy1-S1 sec9-10* cells exited starvation at the restrictive temperature of 36°C, they entered P1, forming SRM domains similar to the wild type (Fig. 1E). Cells then also performed P2, with only a slight delay in SRM domain polarisation as compared to the wild type. In the following hours, cells did not initiate growth confirming that all secretory activity is blocked (Fig. 1E). Such growth arrested cells eventually started dying at around 3hours of SE.

Together, these results indicate that SRM domain formation and polarisation during *de novo* cell polarisation uses a sterol delivery route that neither depends on vesicular traffic via the Golgi apparatus nor on the late secretory machinery.

### Redundant roles of the sterol transporters kes1p and kes3p in *de novo* cell polarisation

An alternative mechanism mediating lipid exchange between different membrane systems occurs via a non-vesicular transport system comprising an evolutionarily conserved family of oxysterol-binding protein (OSBP) and OSBP-related proteins (ORPs) as main transporters ^33^. Fission yeast has six ORP orthologs, none of which is essential (Fig. S2) (Supplementary Materials) ^34^. Three of them, kes1, osh2 and osh3, were previously named based on their sequence homologies to *S.cerevsiae* ORPs, which in case of the latter two includes a distinct protein domain composition (Fig. S2) ^34^. Based on sequence homologies and specific protein features, we named the remaining ORPs kes2p, kes3p and osh8p (for reasoning see Supplementary Materials). To test for a role of these proteins in SE, we analysed all single ORP deletion mutants. All mutants were able to exit starvation and enter exponential growth with near wild type kinetics (Fig. S3A). Similar to *exo70*Δ *sec3-916* cells, we noticed that starved *kes3*Δ cells, and to some extent also *kes1*Δ and *osh8*Δ mutants, had clearly visible small filipin-positive punctae distributed all over the cell periphery. In wild type cells such punctae are barely visible. This suggests a role for ORPs in sterol removal from the plasma membrane in preparation for starvation periods (Fig. S3A). During SE, the punctae persisted, but they seemed immobile as they did not spread to newly formed plasma membrane regions in polar growth sites (Fig. S3B). This indicates that this sterol pool is fixed within the old plasma membrane regions and does not contribute to the formation of new SRM domains.

Since no single ORP is essential, we decided to generate all 15 possible double deletion mutants, one of which we were unable to create (see below). 13 of the remaining did not produce a major phenotype during SE except for some with a moderate delay of P1 and P2 (Fig S3C). Only *kes1*Δ *kes3*Δ double deletion strains showed a significant impact on cell polarisation (Fig. 2A). Similar to the respective single mutants, these cells maintained bright filipin-positive punctae in starvation. In our polarisation ratio measurements, these filipin-positive punctae reduced values of fully polarised cells and they hampered a clear assessment of P1. Qualitative analysis could not fully exclude that, *kes1*Δ *kes3*Δ cells may experience a mild P1 delay (Fig. 2A). P2 in contrast, was substantially delayed and the first clearly polarised cells only became detectable at around 4 hours into SE. The number of polarised cells increased moderately during the following 2 hours. This indicates redundant sterol delivery activity of kes1p and kes3p during P2 of *de novo* cell polarisation.

**Fig. 2:**
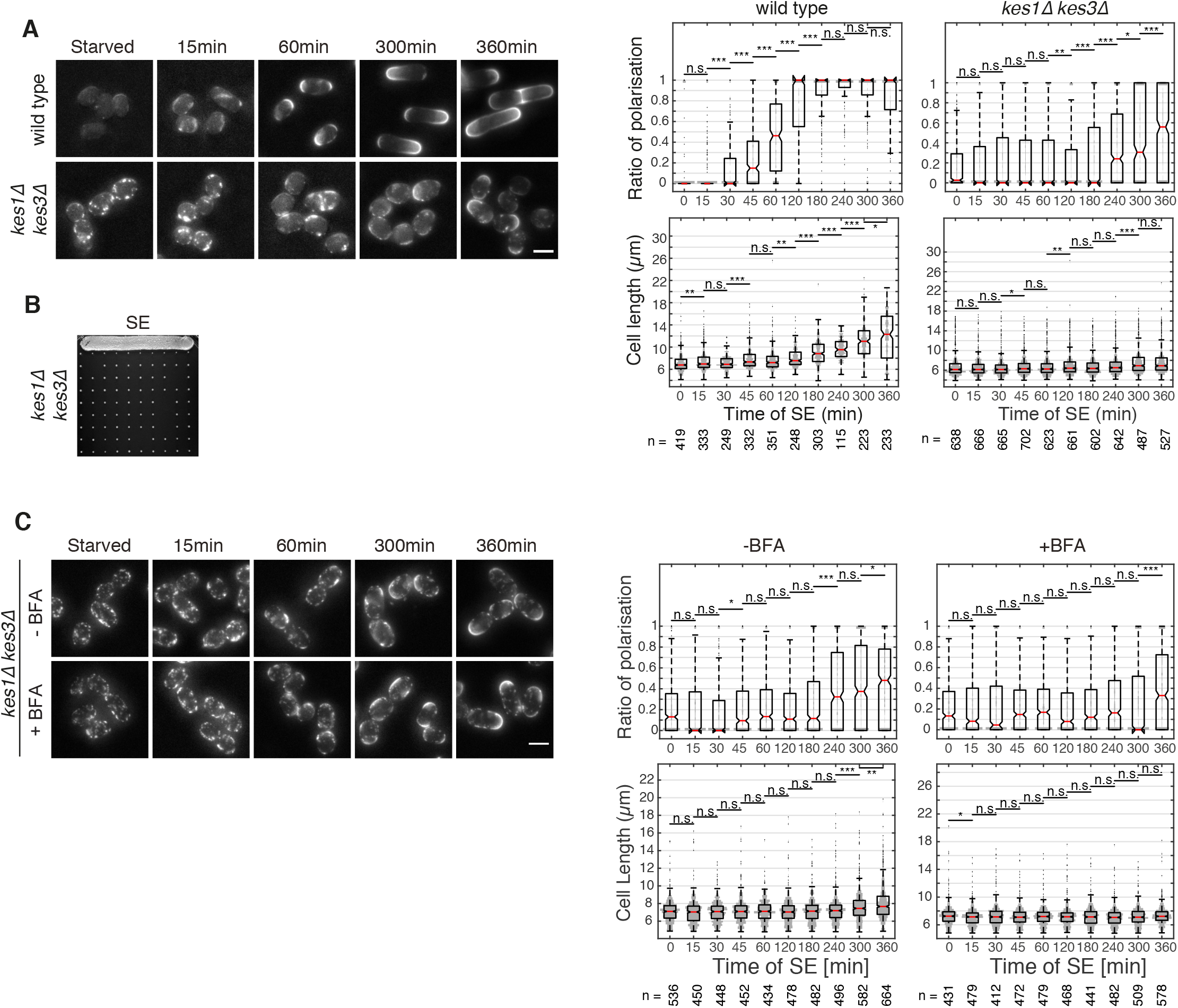
Kes1Δ kes3Δ have a strong polarisation delay. (**A**) SRM domain formation and polarisation during SE, comparing filipin-stained wild type, with *kes1*Δ *kes3*Δ cells. Notched box plots show the polarisation rates and cell lengths at different time points during SE with p-values. (B) Colony forming assay whereby individual starved *kes1*Δ *kes3*Δ cells were arranged on a YE5S plate and allowed to grow into a colony (**C**) SRM domain formation and polarisation during SE, comparing filipin-stained *kes1*Δ *kes3*Δ double mutant cells treated with DMSO (-BFA control) and BFA in DMSO (+BFA). Notched box plots show the polarisation rates at different time points during SE with p-values. Scale bars: 5μm.

Cell length measurements suggested that only a few *kes1*Δ *kes3*Δ cells managed to enter P3 during our 6-hour time-course experiments (Fig. 2A). However, single cell plating of starved *kes1*Δ *kes3*Δ cells showed that most of them eventually initiated growth and were able to form a colony (Fig. 2B). Notably, BFA treatment had almost no additional effect on the strong delay in polarisation of *kes1*Δ *kes3*Δ cells, but, as already shown for wild type cells, it completely blocked P3 (Fig. 2C). This indicates that Golgi apparatus-dependent sterol delivery is not responsible for the cell polarisation capacity remaining in *kes1*Δ *kes3*Δ cells (Fig. 2C).

### Redundantly acting sterol transporters kes1p and osh2p are essential for starvation exit

The only double deletion combination we were unable to generate by crossing was that of *kes1*Δ and *osh2*Δ. We also were not able to do so via the standard homologous recombination-mediated deletion of one gene in a deletion mutant strain of the other. Moreover, our attempts to generate cells expressing either *kes1* or *osh2* under the control of the thiamine repressible nmt81 promoter in *osh2*Δ or *kes1*Δ cells respectively, did not yield a viable strain, notably under non-repressive conditions ^35^. Only a strain that expressed both genes under *nmt81* promoter control (*nmt81:kes1 nmt81:osh2*) was viable. To investigate what causes the sensitivity to the loss of both of these ORPs, we first shut down the *nmt81* promoters by adding thiamine to exponentially growing *nmt81:kes1 nmt81:osh2* cells (Methods). After 4-8 hours - corresponding to 2 division cycles - cells lost most of the filipin staining suggesting that the main sterol delivery routes to the plasma membrane were blocked (Fig. 3A). To test the proliferation capability of such repressed cells we regularly diluted the growing cell population with thiamine-containing medium over a period of 7 days. Throughout this period, sterol delivery remained low, but cells kept proliferating, albeit at a slightly slower rate than wild type cells (Fig. 3B). At around day 2, obvious shape abnormalities, in particular cell diameter fluctuations, appeared (Fig. 3A). Together these results suggest that in growing cells, kes1p and osh2p are not essential but are necessary for efficient sterol delivery. They also show that once polarised, cells can maintain nearly normal polar growth with just small amounts of sterols in the plasma membrane.

**Fig. 3:**
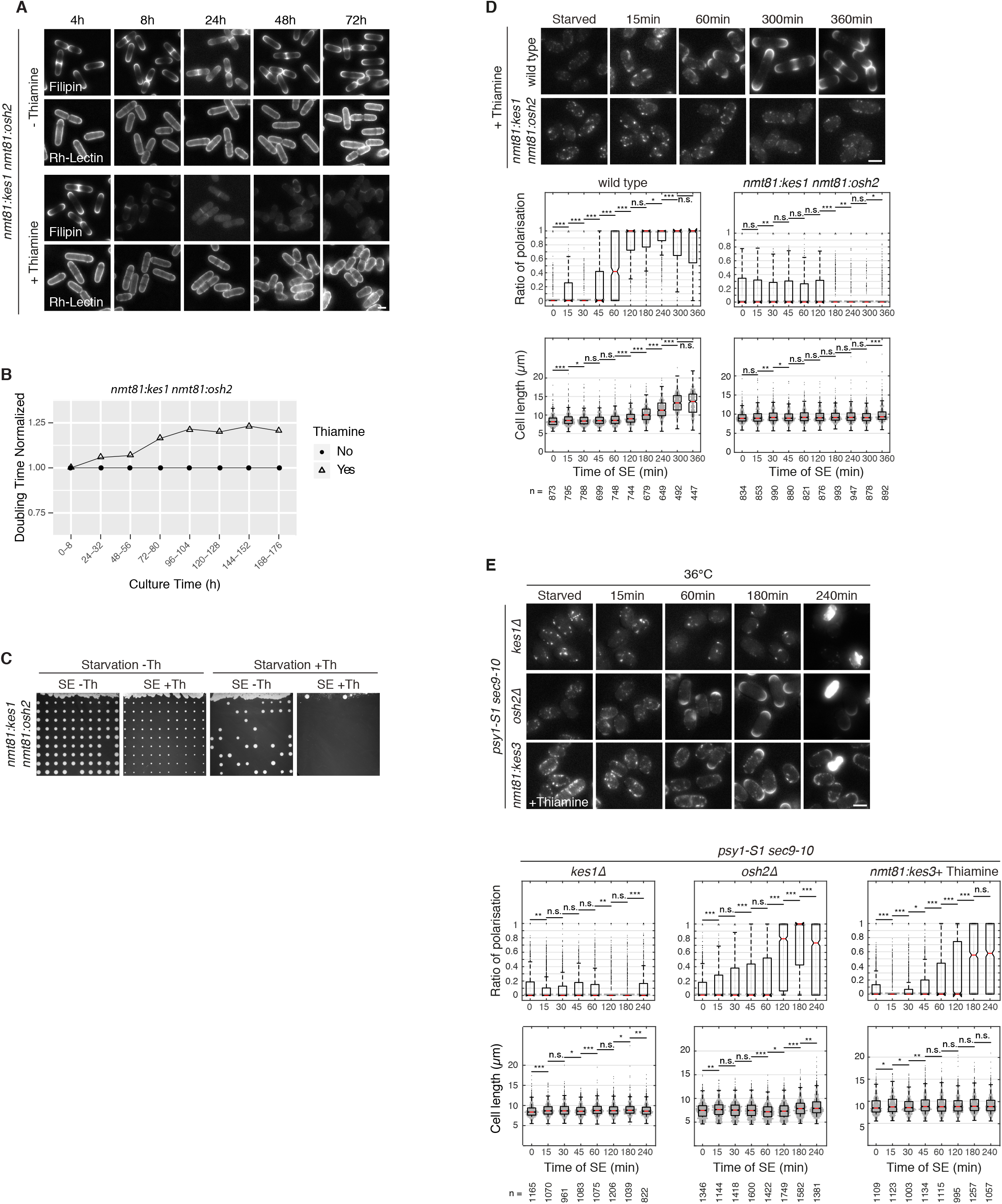
Non-vesicular transport mediates sterol delivery. **(A)** SRM domain maintenance and viability during exponential growth, comparing *nmt81:kes1 nmt81:osh2* cells in medium with or without thiamine. Cells were stained with filipin and Rhodamine-Lectin (for the cell outlines). The contrasted filipin images were equalized to emphasize the reduction of total filipin staining over time. Times after shutoff is indicated in hours (h) **(B)** Graph showing the normalized doubling times of *nmt81:kes1 nmt81:osh2* cells with or without thiamine. The doubling times of cells growing in the absence of thiamine are set to 1.00. **(C)** Colony forming assay whereby individual starved *nmt81:kes1 nmt81:osh2* cells were arranged on EMM2 plates and allowed to grow into a colony. SE-Th and SE +Th indicates the absence or presence of thiamine in the plates respectively. Starvation -Th or +Th indicates the absence or presence of thiamine in the culture medium during starvation entry respectively. (**D**) SRM domain formation and polarisation during SE, comparing filipin-stained wild type, with *nmt81:kes1 nmt81:osh2* cells all in the presence of thiamine. Notched box plots show the polarisation rates and cell lengths at different time points during SE with p-values. (**E**) SRM domain formation and polarisation during SE, comparing filipin-stained wild type, with *nmt81:kes1 nmt81:osh2* cells all in the presence of thiamine. Notched box plots show the polarisation rates and cell lengths at different time points during SE with p-values. Scale bars: 5μm.

We next investigated simultaneous loss of kes1p and osh2p function during *de novo* cell polarisation. If *nmt81:kes1 nmt81:osh2* cells entered starvation with active promoters in the absence of thiamine, followed by SE in the presence of thiamine, cells exited starvation and formed colonies in single cell growth assays. However, the colonies were very small as compared to those of control cells lacking nmt81 suppression (Fig. 3C). Since thiamine-mediated suppression of gene expression only leads to elimination of protein with a certain time delay, it is possible that under these experimental conditions, sufficient protein remains to allow SE.

To ensure that protein levels were maximally depleted in quiescent cells prior to SE induction, we repressed the nmt81 promoters by adding thiamine already when moving *nmt81-kes1 nmt81-osh2* cells into starvation medium (Methods). SE was then initiated and observed under continued thiamine repression. Strikingly, the cells showed no sterol delivery to the plasma membrane under these conditions and they remained fully arrested (Fig. 3D). This reveals an essential, redundant role of kes1p and osh2p during the initial delivery of sterols to the plasma membrane in P1 of *de novo* cell polarisation. Single cell plating of such starved *nmt81:kes1 nmt81:osh2* cells showed that less than 1% of them managed to exit starvation in the presence of thiamine. In contrast, around 40% of *nmt81:kes1 nmt81:osh2* cells exited starvation in a rescue experiment, where no thiamine was added to the medium triggering SE (Fig. 3C). Notably, reversing thiamine repression is a very slow process and is suggested to take 10-15 hours in proliferating cells. This obviously includes dilution of promotor-bound thiamine during every S-phase ^36^. The fact that no S-phase and thus no such dilution occurs in our rescue experiment provides a likely explanation for the remaining 60% cell lethality, which most likely is due to persistent thiamine repressive activity. This is consistent with the previously observed lethality of cells trying to exit starvation with drug-mediated inhibition of sterol production ^17^.

### Kes1p, but not osh2p or kes3p, directly delivers sterols to the plasma membrane

The uncovered redundancy between kes1p and osh2p in P1, may occur on a functional level where kes1p and osh2p substitute for each other’s function. Alternatively, the two proteins are part of two independent sterol delivery pathways. The latter scenario raises the possibility that one of the two pathways includes vesicular transport after all, which in our secretory mutants would have been masked by the other pathway. To investigate this further, we blocked vesicle fusion with the plasma membrane in *psy1-S1 sec9-10* mutants in addition carrying a *kes1*Δ or *osh2*Δ deletion. When following SE of these cells at the restrictive temperature *psy1-S1 sec9-10 osh2*Δ cells showed almost normal cell polarisation behaviour except that P2 entry was somewhat delayed (Fig. 3E). Cells subsequently were unable to enter P3 and eventually died similar to *psy1-S1 sec9-10* cells. This suggests that kes1p efficiently delivers sterols directly to the plasma membrane during P1, without the involvement of the late secretory pathway. In contrast, *psy1-S1 sec9-10 kes1*Δ cells exiting starvation, showed no sign of sterol delivery and SRM domain formation whatsoever (Fig. 3E). This indicates that unlike kes1p, osh2p activity requires the fusion of vesicles with the plasma membrane, placing osh2p function somewhere along a vesicular sterol delivery route. These results reveal that during P1 of SE, *de novo* synthesised sterols are delivered to the plasma membrane via two independent, redundantly acting routes that differ in their requirement for vesicular sterol transport.

The strong synthetic effect in *de novo* cell polarisation of the *kes1*Δ and *kes3*Δ deletions described above, raises the same question of redundancy as with kes1p and osh2p. Thus, from the *psy1-S1 sec9-10 kes1*Δ cells we can similarly conclude that also the kes3p function, which provides nearly normal cell polarisation in *kes1*Δ cells, depends on SNARE-mediated vesicle fusion with the plasma membrane. Interestingly, we failed to delete *kes3* in *psy1-S1 sec9-10* cells via standard homologous recombination. We therefore introduced the *nmt81* promoter in front of the endogenous *kes3* gene in *psy1-S1 sec9-10* cells, and repressed it by adding thiamine during starvation entry as before with the *nmt81:kes1 nmt81:osh2* cells. As in *kes3*Δ single mutants, filipin positive punctae remained at the plasma membrane in the starved cells confirming efficient kes3p depletion. When exiting starvation at the restrictive temperature, *nmt81:kes3 psy1-S1 sec9-10* cells formed and polarised SRM domains similar to *psy1-S1 sec9-10 osh2*Δ cells confirming kes1p acting independent of the late secretory pathway (Fig. 3E). In summary, our results show that kes1p directly delivers sterols to the plasma membrane whereas the redundant activities of osh2p and kes3p provide a non-vesicular sterol transport step in a vesicular sterol delivery pathway.

### Kes1p provides the main polarising activity in P2

Cdc42p, was previously shown to be able to spontaneously polarise budding yeast cells ^37^. Yet, in *de novo* polarising fission yeast cells, cdc42p was found to be dispensable and accordingly its activity was detectable only after cells had repositioned their SRM domains to the cell poles in P2 ^17^. We hypothesised that this late onset cdc42p activity may be responsible for the rescue of polarisation in *kes1*Δ *kes3*Δ cells. To investigate this possibility, we individually combined the *kes1*Δ and *kes3*Δ deletions with the temperature sensitive *cdc42* allele *cdc42-3* ^38^. Already at permissive temperature *cdc42-3* causes a particularly strong loss-of-function phenotype that produces cells with a pronounced spherical morphology (Fig. 4A). Notably, when exiting starvation at the restrictive temperature (36°C) such rounded cells were struggling to produce the bipolar microtubule arrays known to be required for polarised tea1p deposition (Fig. 4B). Consequently, the most affected *cdc42-3* cells showed a delay in SRM domain polarisation during P2 (Fig. 4C). This delay is expected as we had previously shown that SRM domain polarisation in P2 fully depends on polar tea1p deposition ^17^. Yet, after 3 hours at the restrictive temperature, most cells displayed the typical SRM caps at both cell poles suggesting an otherwise functional cell polarisation process (Fig. 4C). At this stage some cells had small, centrally located SRM domains. Normally, this would suggest incomplete SRM domain removal from the cell middle. However, at later time points, the number of such centrally located domains considerably increased again suggesting that once polar SRM domain organisation was established, cells were unable to maintain this state. Supporting this, *cdc42-3* cells, at around this time, initiated growth also at the restrictive temperature. However, growth was not polar since at 6 hours of SE cells had similarly enlarged in length and width (Fig. 4C). This is consistent with the previously proposed essential role of cdc42p in maintaining polar growth rather than in establishing it.

**Fig. 4:**
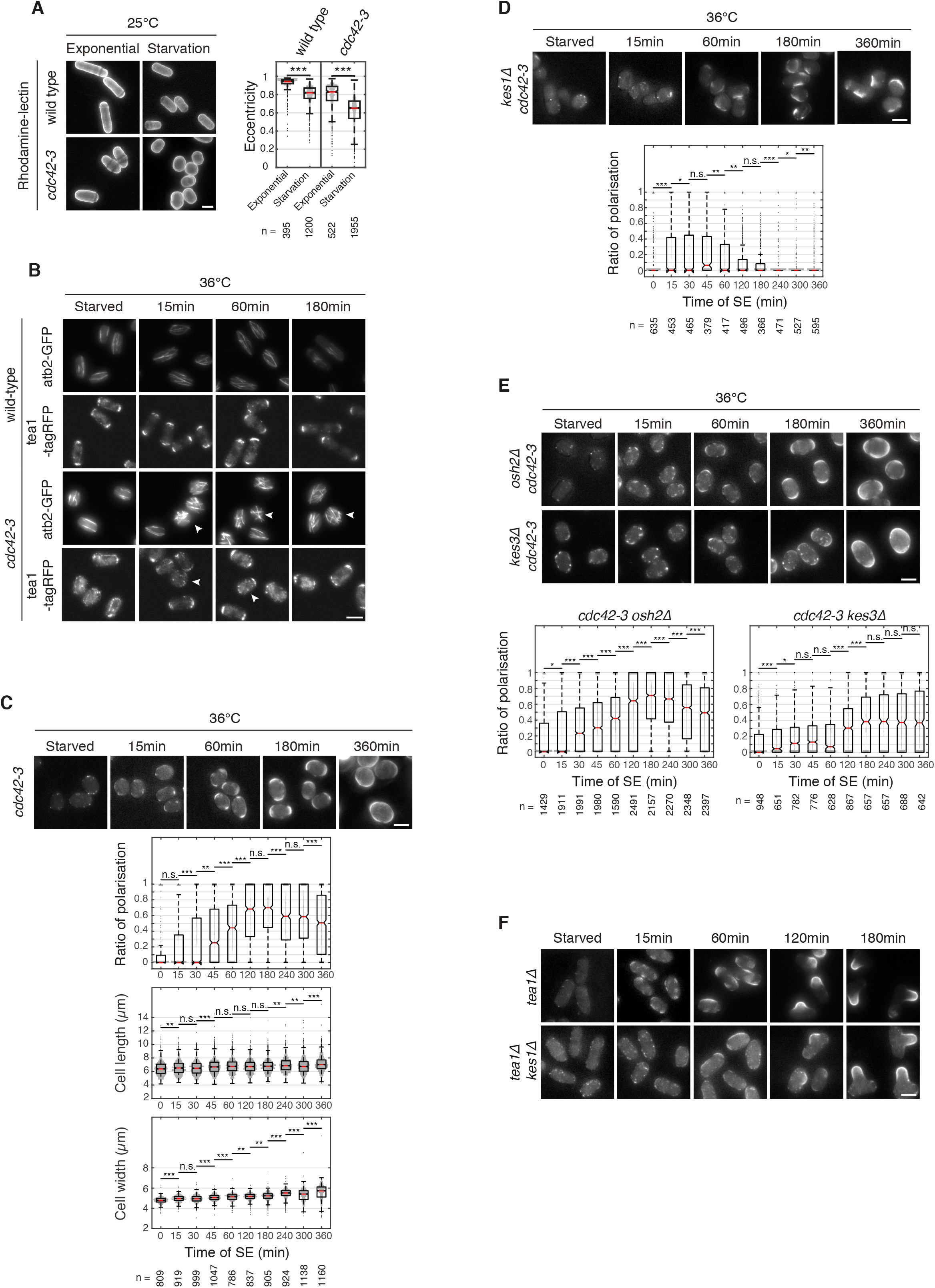
Cdc42p/Kes3p provide a redundant polarisation pathway. (**A**) Qualitative and quantitative comparison of Rhodamine-Lectin-stained wild type and *cdc42-3* mutants grown at the permissive temperature (25°C) and imaged in exponential growth or in starvation after 3 days of culturing. Notched box plots show the distribution of cellular shape eccentricity (Supplementary Materials). (**B**) Comparison of microtubule organization and tea1p localization in wild type and *cdc42-3* mutant cells expressing GFP-tagged α2-tubulin (atb2-GFP) or RFP-tagged tea1p (tea1-tagRFP) at different time points during SE at the restrictive temperature (36°C). (**C**) SRM domain formation and polarisation during SE at the restrictive temperature (36°C), showing filipin-stained *cdc42-3* cells. Notched box plots show the polarisation rates at different time points during SE with p-values. (**D**) SRM domain formation and polarisation of filipin-stained *cdc42-3 kes1*Δ cells at different time-points during SE at the restrictive temperature (36°C). Notched box plots show the polarisation rates at different time points during SE with p-values. (**E**) SRM formation and polarisation of filipin-stained *cdc42-3 kes3*Δ and *cdc42-3 osh2*Δ cells at different time-points during SE at the restrictive temperature (36°C). Notched box plots show the polarisation rates at different time points during SE with p-values. (**F**) SRM domain formation and polarisation during SE, comparing filipin-stained *tea1*Δ and *tea1Δ kes1*Δ cells Scale bars: 5μm.

This was very different for *kes1*Δ *cdc42-3* double mutant cells exiting starvation at the restrictive temperature. Similar to *cdc42-3* single mutants, these cells entered P1 and formed one or two randomly distributed SRM domains (Fig. 4D). However, *kes1Δ cdc42-3* cells were unable to enter P2 and even after 6 hours, polar SRM domains were rare, as expected for a random distribution. This indicates that in *kes1*Δ mutants, SRM domain polarisation becomes cdc42p dependent. Based on these findings, we conclude that the osh2p- and kes3p-dependent sterol delivery routes rescuing loss of kes1p in P1 and P2 respectively, are under cdc42p control. Consistently, *osh2*Δ *cdc42-3* and *kes3*Δ *cdc42-3* double mutants formed and polarised their SRM domains similar to the *cdc42-3* single mutant (Fig. 4E). These results, together with the previously reported late-P2 onset of cdc42p activity, suggest that in wild type cells, the strictly non-vesicular kes1p activity provides the main sterol delivery route driving microtubule/tea1p-mediated SRM domain polarisation in P2.

Since cdc42p polarising activity was mainly studied in exponentially growing cells, in which the tea1p-independent polarity inheritance pathway dominates, this raises the possibility that after all cdc42p may not be under direct control of the microtubule/tea1p system as is commonly assumed. To explore whether the accurate polar SRM domain positioning in *kes1*Δ cells by the cdc42p/kes3p activity requires tea1p, we followed *de novo* cell polarisation of *kes1*Δ cells in which *tea1* was additionally deleted (*kes1*Δ *tea1*Δ). The resulting cells displayed a typical *tea1*Δ phenotype with cells harbouring a single, randomly positioned growth site (Fig. 4F) ^17^. These data confirm that the positioning of cdc42p activity is under the control of the tea1/microtubule system also during *de novo* cell polarisation in cells lacking the preferred kes1p sterol delivery route.

### Sterol transport requires PI4P

Non-vesicular sterol trafficking by some ORPs was proposed to involve counter-flow transport of sterols and phosphatidylinositol 4-phosphate (PI4P) ^39^. To test a requirement for acute synthesis of PI4P during cell polarisation, we analysed SE of cells expressing an engineered version of pik1p, the kinase producing PI4P, which carries a temperature sensitive protein degradation module (*pik1-td*) ^40^. Similar to starved *kes1*Δ and *kes3*Δ cells, starved *pik1-td* cells showed filipin-positive punctae distributed all over the plasma membrane suggesting that already the efficient sterol removal during starvation entry is PI4P-dependent (Fig. 5A). When exiting starvation at the restrictive temperature, the very first SRM domains became clearly visible only after about 3 hours, suggesting strong impairment of P1 (Fig. 5A). In the following hour, the number of cells with SRM domains increased only minimally. Within 6 hours, some cells accumulated sterols in a single, prominent domain that could also be situated at a cell pole. The slight increase in the polarisation ratio suggested that some of the cells had managed to go through P2. However, with a certain probability, P1 cells with a single SRM domain will also have it situated at a cell pole. Due to the low number of such cells we could not determine to what extent pik1p is required also to perform P2. Clear was that *pik1-td* cells could not enter P3 and initiate growth at the restrictive temperature. Eventually, these cells died.

**Fig. 5:**
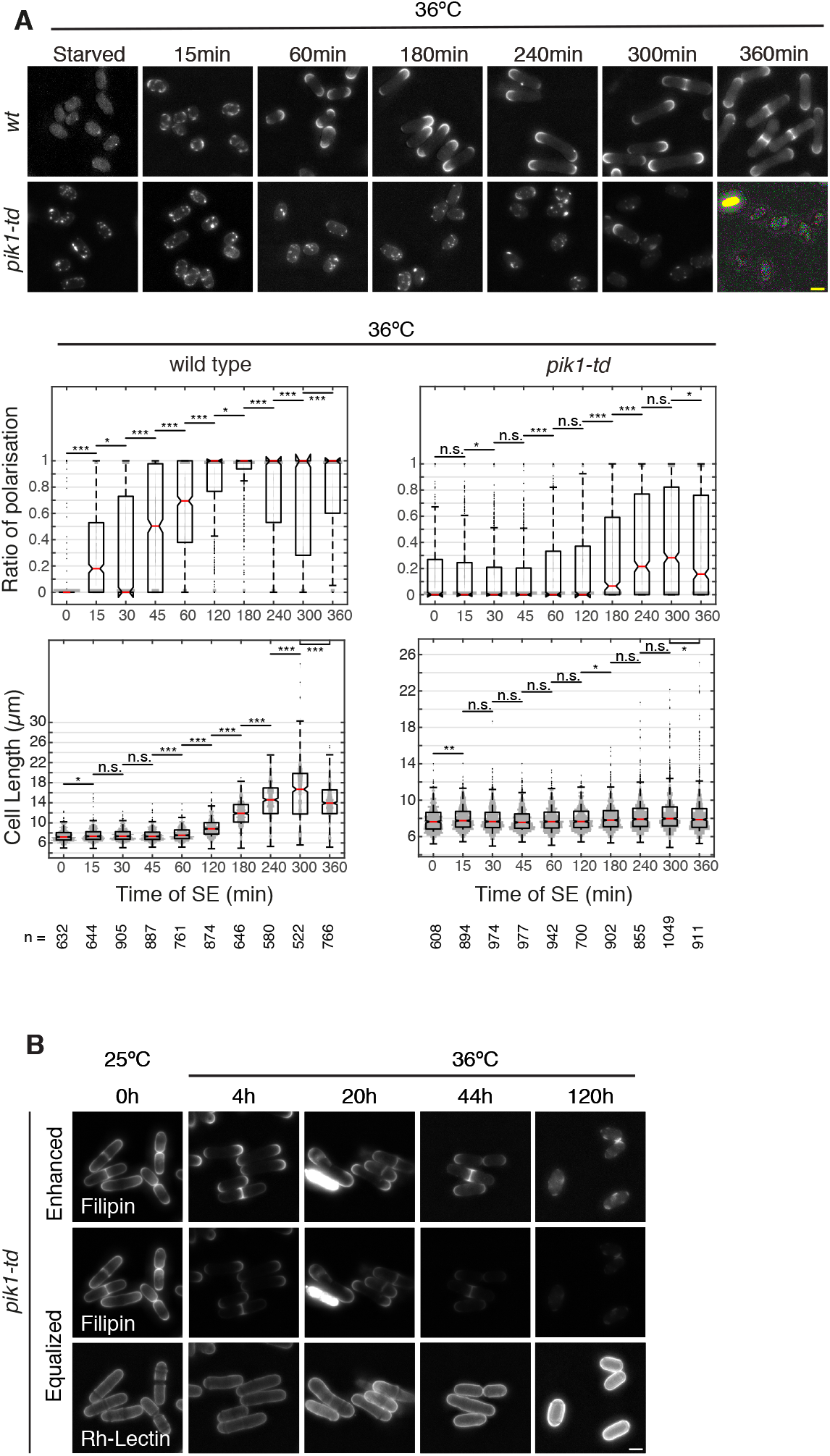
PI4P drives sterol delivery to the PM. (**A**) SRM domain formation and polarisation during SE, comparing filipin-stained wild type, with *pik1-td* cells at the restrictive temperature (36°C). Notched box plots show the polarisation rates and cell lengths at different time points during SE with p-values. **(B)** Time course showing exponentially growing *pik1-td* cells after a shift to restrictive temperature (36°C). Cells were stained with filipin and Rhodamine-Lectin (for the cell outlines). Two differently contrasted images of the filipin stained cells are shown to emphasize the loss of SRM polarity (Enhanced) or the reduction of total filipin staining over time (Equalized). Time indicated in hours (h) after the shift to 36°C. Scale bars: 5μm.

We conclude that phosphoinositides are crucial for sterol delivery to the plasma membrane in *de novo* polarising cells. Notably, our results show that this is the case for both the t-SNARE-dependent and the t-SNARE-independent pathway acting during P1. In addition, this also applies to sterol-removal from the plasma membrane in cells entering starvation.

In agreement with a role for PI4P in ORP-mediated sterol delivery to the plasma membrane, we observed that *pik1-td* cells growing at the permissive temperature, when shifted to the restrictive temperature, developed a phenotype that was very similar to that of exponentially growing *nmt81:kes1 nmt81:osh2* cells under repressed conditions: Already after 4 hours *pik1-td* cells lost most of the filipin staining at the cell poles and the septa and after 5 days of continued culturing very little detectable signal remained (Fig. 5B).

### Tea1p does not regulate localisation of kes1p, kes3p or pik1p

The random positioning of the newly formed SRM domains in P1 and their tea1p-controlled polarisation in P2, could potentially correlate with a corresponding localisation of the responsible ORPs and/or pik1p. In particular in P2, tea1p might polarise SRM domains by positioning kes1p or the production of PI4P. To test this possibility, we monitored GFP-tagged protein versions during SE. For kes1p, kes3p and osh2p we introduced the monomeric mGFP gene sequence at the c-terminus of the respective endogenous gene loci ^41^. For pik1p, we used a previously published version that was n-terminally tagged with GFP(S65T) ^40^. None of the proteins showed a distinct localisation at the plasma membrane at any phase of SE. All proteins evenly stained the entire cytoplasm. In GFP-pik1p and kes3p-mGFP cells, significant signal additionally accumulated in the nucleus (Fig. 6A, B). Furthermore, in osh2p-mGFP cells 1-2 sub-cellular spots were visible at 3hours of SE, which also were detectable in exponentially growing cells. We conclude that SRM domain positioning during *de novo* cell polarisation is not controlled by regulation of ORP or pik1p localisation.

**Fig. 6:**
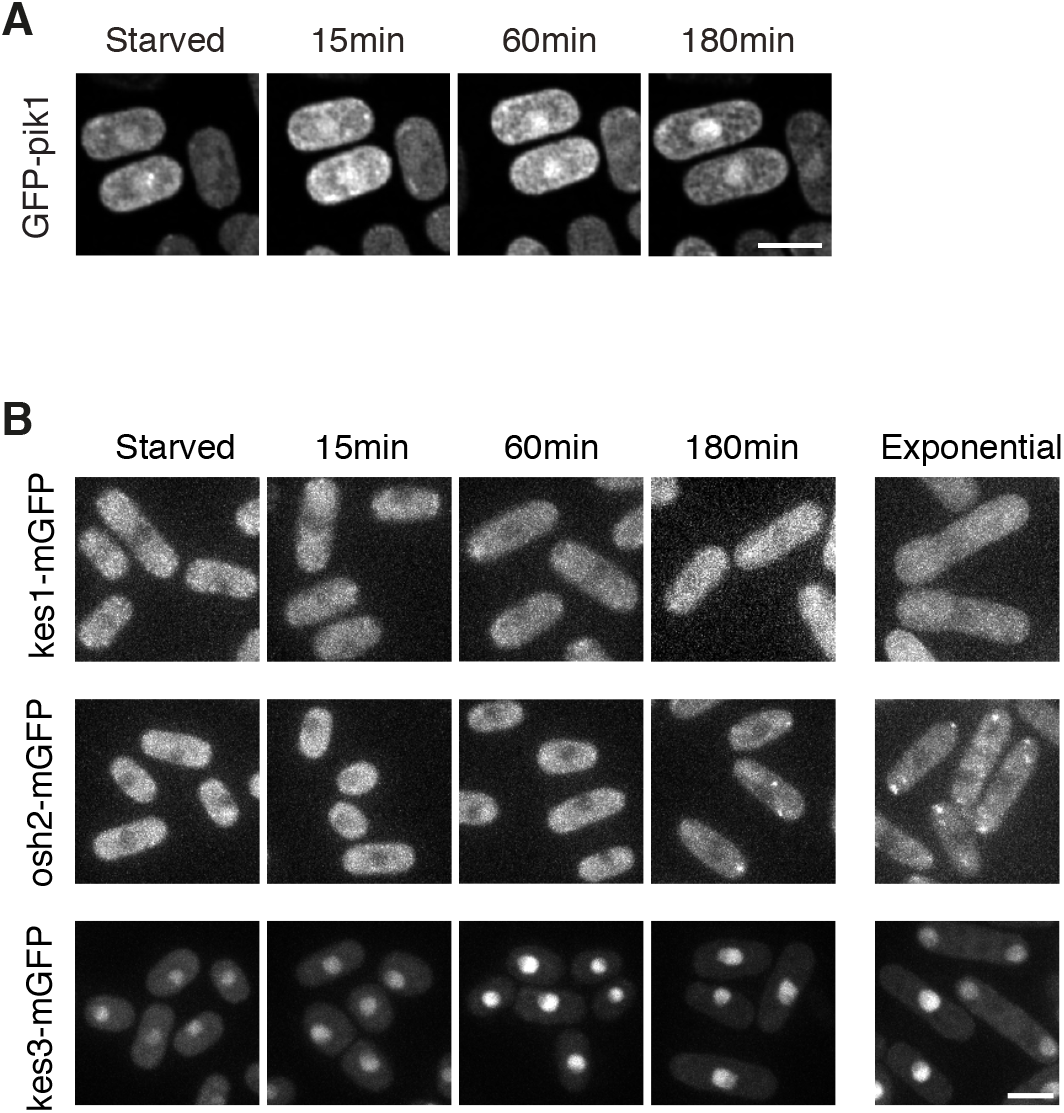
Localizing pik1p, kes1p, osh2p and kes3p. **(A)** localization of GFP-pik1p during SE. **(B)** localization of kes1p-mGFP, osh2p-mGFP, and kes3p-mGFP during SE and in exponential phase. Scale bars: 5μm.

Notably, we also tagged the kes1p and kes3p proteins with GFP(S65T), which unlike mGFP is sticky and can mediate protein aggregation. With this GFP variant, we indeed detected more distinct localisation during SE. Both, kes1p-GFP(S65T) and kes3p-GFP(S65T), started showing faint signal all over the cell periphery suggesting global accumulation near the plasma membrane. Consistent with its early role, kes1p-GFP(S65T) did so after 15minutes, while peripheral kes3p-GFP(S65T) only became visible at around 45 minutes - fitting well with its dependence on cdc42p activation. Kes1p-GFP accumulated in additional sub-cellular structures at around the time when cells entered P3 and initiated growth. This could relate to additional functions in growth homeostasis, as suggested for the ORP orthologs in budding yeast (Fig. S4A) ^42,43^.

## Discussion

OSBP/ORP family proteins have been extensively studied, making clear that non-vesicular sterol transfer between various membranes is an important activity affecting numerous cellular activities. A key cell biological function for this transport activity was assigned to sterol homeostasis providing an asymmetric distribution of sterols in the cellular membrane system ^44^. Besides, some ORPs were shown to contribute to the tethering of membrane contact sites, to homeostasis of other lipids and they were implicated in the regulation of vesicular trafficking, including COPII-mediated vesicle transport and polarised exocytosis ^42–51^. In particular in budding yeast, a link to the maintenance of Cdc42p-controlled cell polarity was described ^52^. However, this activity is not specific to a particular ORP. Accordingly, budding yeast cells are viable when all but any one of its seven ORPs are deleted, showing high redundancy amongst these proteins ^53^. We now describe a vital role for only two out of the six fission yeast ORPs during *de novo* cell polarisation. This process was previously shown to depend on the delivery of *de novo* synthesised sterols and its regulation by the microtubule/tea1p polarity system ^17^. We genetically dissected two redundant, but differing sterol delivery pathways, which mediate the formation of the first, “priming” SRM domains in P1 of SE (Fig. 7A). One pathway involves kes1p, which delivers sterols directly to the plasma membrane. The other pathway combines osh2p function, with vesicular transport, requiring vesicle fusion at the plasma membrane. Kes1p and osh2p thus are non-redundant with respect to their molecular function. Notably, both pathways function independent of the Golgi apparatus (Fig. S5A). Together, the two delivery routes are absolutely essential not only for any further step of cell polarisation and growth initiation but also for cell survival. This shows that kes1p and osh2p safeguard a vital function that cannot be complemented by any of the other four fission yeast orthologs.

**Figure 7:**
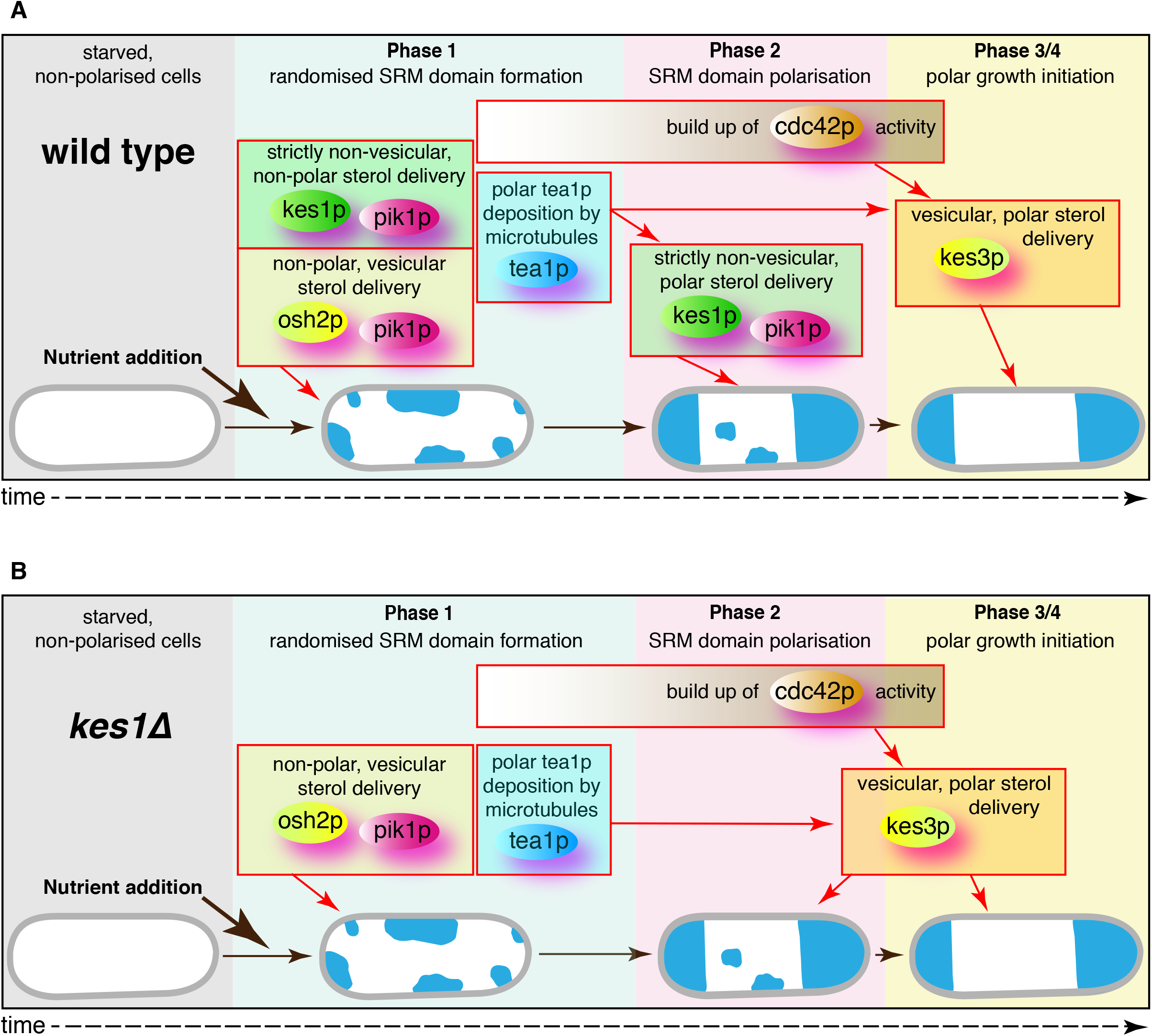
Cartoon of *de novo* cell polarisation. The cartoon correlates the requirement of the most relevant proteins in this study with the phases of SE proteins shows the sequence of events we propose for microtubule/tea1p-guided SRM domain formation and polarisation during SE (**A**) in wild type cells and (**B**) in *kes1*Δ cells. Red arrows indicate the proposed regulatory hierarchy, without implying direct interaction.

In P2, wild type cells rely on the kes1p pathway, while the osh2p pathway does not play a detectable role. This we can conclude, because in the absence of cdc42p activity, cells have to produce kes1p to proceed through P2, while the osh2p pathway is dispensable. Cdc42p in turn, provides a redundant activity that rescues P2 in cells lacking kes1p function. However, in wild type cells it was previously shown that cdc42p activity only becomes detectable at the end of P2, indicating that normally the kes1p delivery route drives most of this decisive phase of microtubule-controlled cell polarisation ^17^.

Our results obtained with cells lacking the kes1p pathway, reveal that cdc42p controls a third sterol delivery route, which safeguards SRM domain polarisation following P1. A third route explains the moderate delay of SRM domain polarisation observed in *kes1*Δ cells. This route comes into effect as soon as cells have attained sufficient cdc42p activity explaining why SRM domain polarisation in *kes1*Δ cells is only moderately delayed. It additionally provides a possible mechanism underlying so far unexplained aspects of our previous work: In Makushok et al. we reported that the onset of P3, occurred shortly before SRM domain polarisation had completed ^17^. Presumably, this marks the moment when the levels of cdc42p activity at one of the enlarging, polar SRM domains has reached the critical level for triggering growth. This would simultaneously enforce sterol delivery to the growing cell pole. Consistently, this moment was reported to initiate a temporary weakening of the SRM domain at the opposite cell pole, suggesting a transient competition for a given sterol pool ^17^.

These conclusions raise the question about the nature of this cdc42p-dependent, third sterol delivery route. The profound P2 delay in *kes1*Δ *kes3*Δ cells, indicates that kes3p activity is critical. We can think of two possible mechanisms: In the first, kes3p depicts a cdc42p-controlled sterol delivery pathway that functions independent of kes1p and osh2p (Fig. 7B). In the second, kes3p and osh2p are part of the same sterol delivery pathway, in which they mediate independent non-vesicular sterol transport steps. Notably, this osh2p activity would differ from its cdc42p-independent activity in P1. Furthermore, the fact that P1 and cell viability in SE critically depend on the presence of either osh2p or kes1p, while *osh2*Δ *kes3*Δ cells are fully viable and also *kes1*Δ *kes3*Δ cells still make it all the way to form growing colonies, strongly support the scenario in which the cdc42p-controlled kes3p pathway acts independently of kes1p and osh2p.

Regardless of the exact molecular composition of the different pathways, a remaining question is how *kes1*Δ *kes3*Δ cells eventually manage to polarise. Given the high degree of redundancy amongst ORPs in budding yeast, it seems plausible that one or more of the remaining ORPs substitute, although poorly, for kes3p function in the double mutant ^53^. The less likely second possibility is that *kes1*Δ *kes3*Δ cells can make use of yet another sterol delivery route. This route would have to function independent of the Golgi apparatus, which can be eliminated in *kes1*Δ *kes3*Δ cells without much effect. Obviously, this route would not be able to compensate for the loss of kes1p and osh2p activity in P1 and would be controlled by cdc42p. A possible scenario is the recycling of the sterols that were delivered to the plasma membrane during P1. Such a mechanism was shown to be crucial during polar growth in mating budding yeast cells ^54,55^.

Altogether, our results reveal a key role for kes1p-mediated sterol delivery in *de novo* polarising wild type cells that is independent of SNARE-dependent vesicle fusion at the plasma membrane. The most plausible mechanism is that kes1p transports the newly synthesised sterols directly from their production site, the ER, to the plasma membrane. In fission yeast, these two membranes are closely aligned throughout the cell circumference, also in starved cells, which certainly facilitates such a mechanism ^24,56^. Yet, at this stage we cannot fully exclude the involvement of an unidentified membrane organelle intermediate, acting prior to kes1p in a multi-step pathway.

Our findings provide a new twist to the old question as to how the microtubule/tea1p system controls *de novo* cell polarisation. One now needs to understand how polarly deposited tea1p governs the switch from randomised kes1p and osh2p activity in P1, to polar kes1p activity in P2. Our results exclude direct regulation of kes1p localisation. Previous findings with a live marker of SRM domains, render a mechanism, in which sterols are randomly delivered and then laterally sorted within the plasma membrane, unlikely ^17^. Therefore, localised regulation of kes1p activity is the most probable scenario. One possibility is that localised availability of phosphoinositides plays a role. Previous work had suggested that some ORPs, exchange the bound sterol with a PI4P when arriving at the sterol acceptor membrane. PI4P is then transported back to the donor membrane ^39^. Consistently, impaired PI4P production during SE, nearly blocked SRM domain formation and polarisation. The few escaping cells most likely reflect mutant leakiness, as found in most experiments using temperature sensitive protein interference. However, at this stage we cannot rule out indirect effects on sterol delivery in pik1p mutants, including defective sterol production.

We here identify non-vesicular sterol transport as essential component of *de novo* cell polarisation and growth initiation, corroborating the absolutely central role of SRM domains in cell polarisation postulated by Makushok et al. ^17^. The role of the microtubule/tea1p system therefore shifts away from direct regulation of the proteins constituting the polarisome to proteins regulating non-vesicular sterol transport ^57^. This is also the case for the alternative, cdc42p-controlled delivery route that becomes essential in the absence of kes1p activity. In such a scenario, the role of cdc42p in growth polarity remains essential, but in wild type cells it would mainly serve to enforce and/or maintain a polarised state rather than being central to establishing it. Consistently, fission yeast cdc42p alone cannot polarise a starved cell *de novo* without the priming SRM domains first having formed in the plasma membrane during P1. Notably, the classical experiment showing *de novo* polarising activity of constitutively active Cdc42 in budding yeast mutants, was performed with cells in which sterols were already present in the plasma membrane ^10^. All our findings thus are still fully in line with an evolutionarily conserved mechanism mediating the first steps of cell polarisation also in higher eukaryotes.

## Supporting information

Analysis_Software_Details

## Acknowledgments

We thank Darren Gilmour, and Angela Lloyd for critical reading of the manuscript. We are grateful to Yauhen Yakimovic and Maria B. Heimlicher for advice on image segmentation. We thank Grégory Paul for advice on statistical issues. This work was funded by a grant from the Swiss National Science Foundation to DB.

## Data availability statement

The manuscript does not contain publicly available datasets.

Figures 1B-E, Figures 2-6, Fig S1 and Figures S3-S5 are associated with raw data

There are no restrictions on data availability. Raw imaging data and yeast strains can be obtained from S.M.H or D.B upon request.

## Code availability statement

The code for our analysis software is available Bitbucket - https://bitbucket.org/Dreher/yeast-border-trace/.

## Supplementary Materials

### List of supplementary Materials

**Fig. S1:**
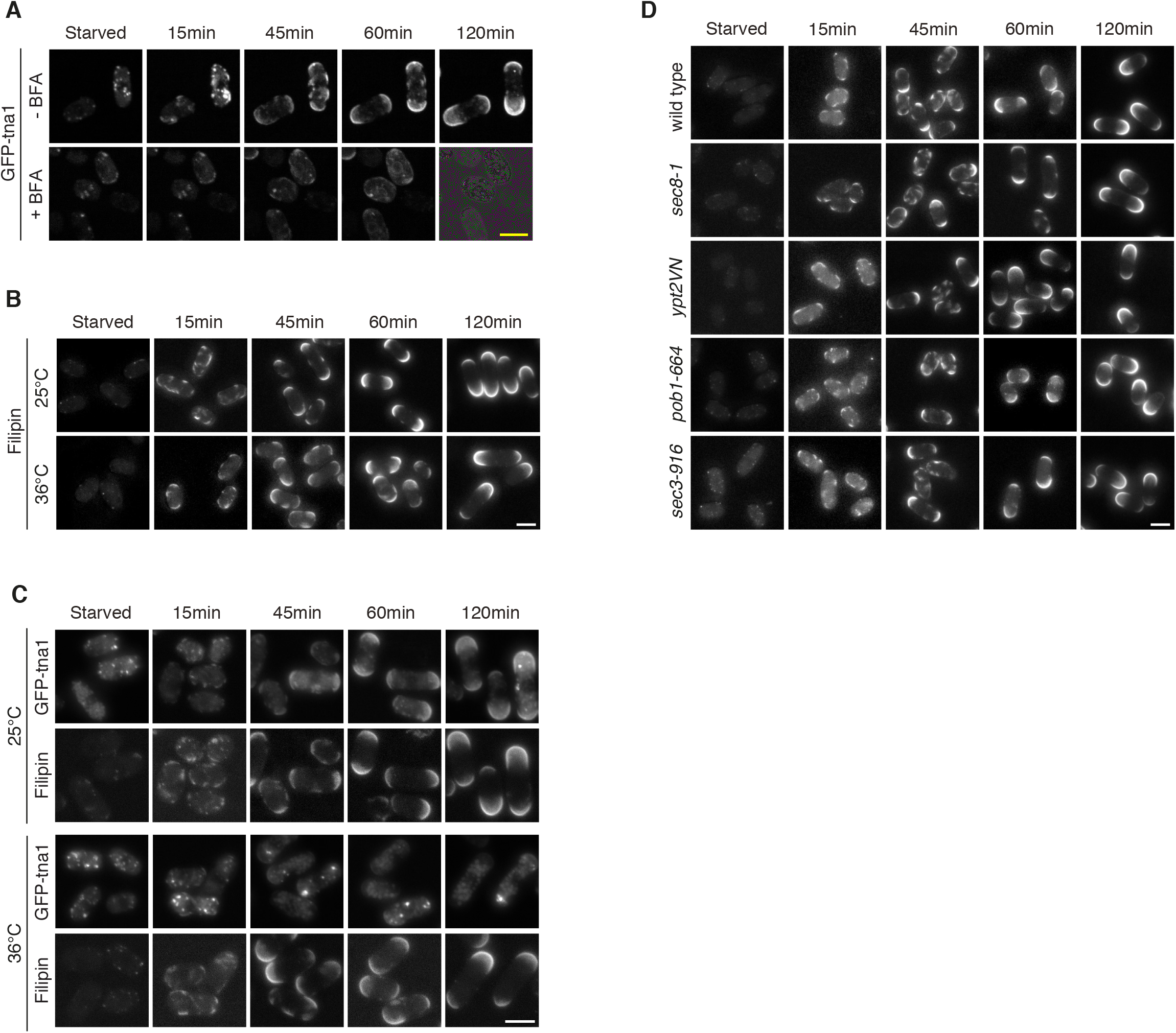
Sterol delivery occurs independent of an intact Golgi apparatus. (**A**) Localisation of GFP-tagged tna1p during SE of wild type cells treated with DMSO (-BFA control) and BFA in DMSO (+BFA). (**B**) SRM domain formation and polarisation during SE, comparing filipin-stained wild type cells at 25°C and 36°C. (**C**) Qualitative comparison of localization of GFP-tagged tna1p during SE of wild type cells at 25°C and 36°C. (**D**) SRM domain formation and polarisation during SE at the restrictive temperature (36°C), comparing filipin-stained wild type cells with different mutants of vesicular secretion. Scale bars: 5μm.

**Fig. S2:**
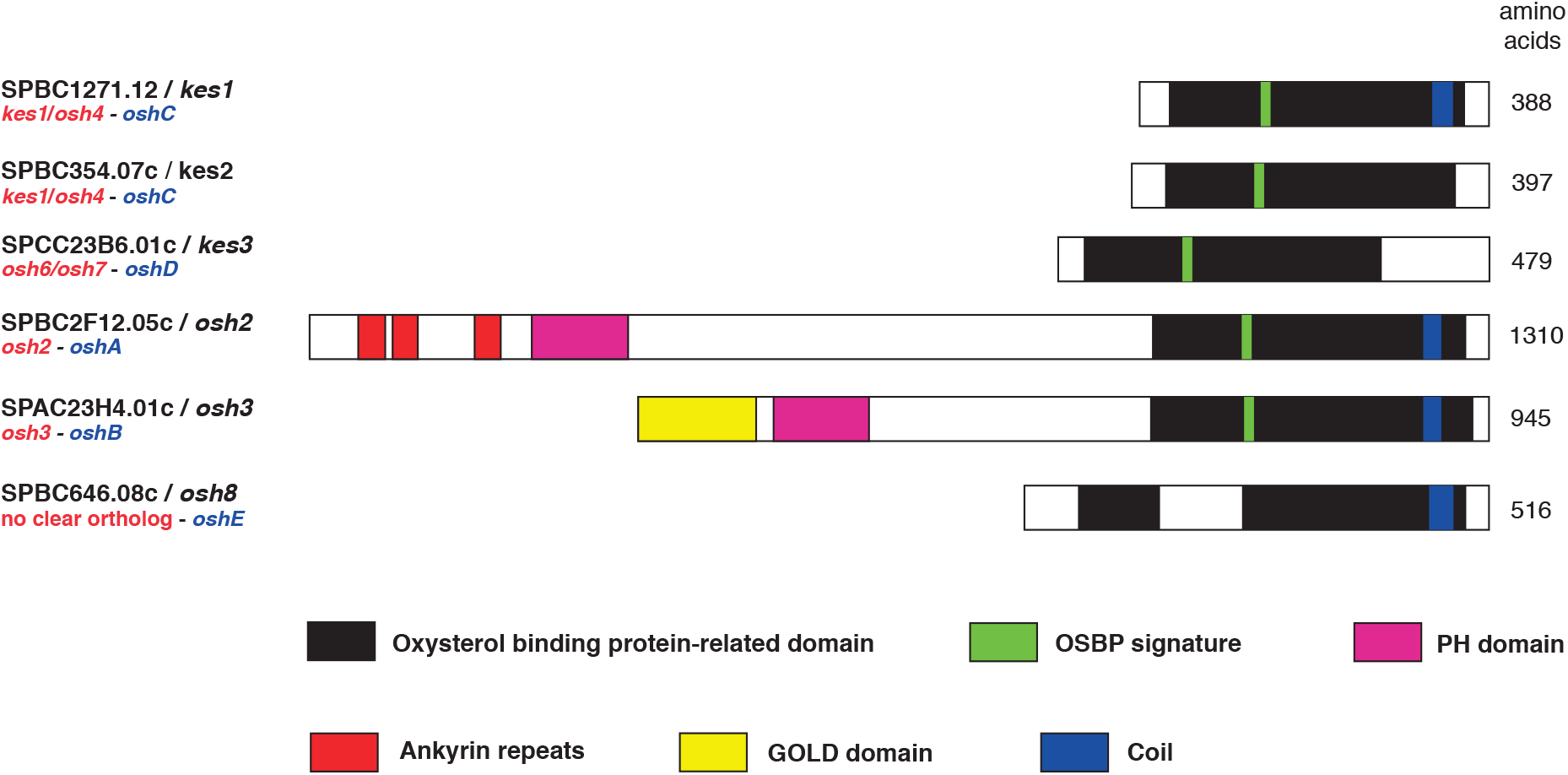
*S. pombe* ORPs and their protein domains. Protein domains as annotated in Pombase are highlighted in colour (*21*). Where missing, gene names were assigned based on closest overall protein sequence homology to ORPs in budding yeast (names in red) and *Aspergillus nidulans* (names in blue).

**Fig. S3:**
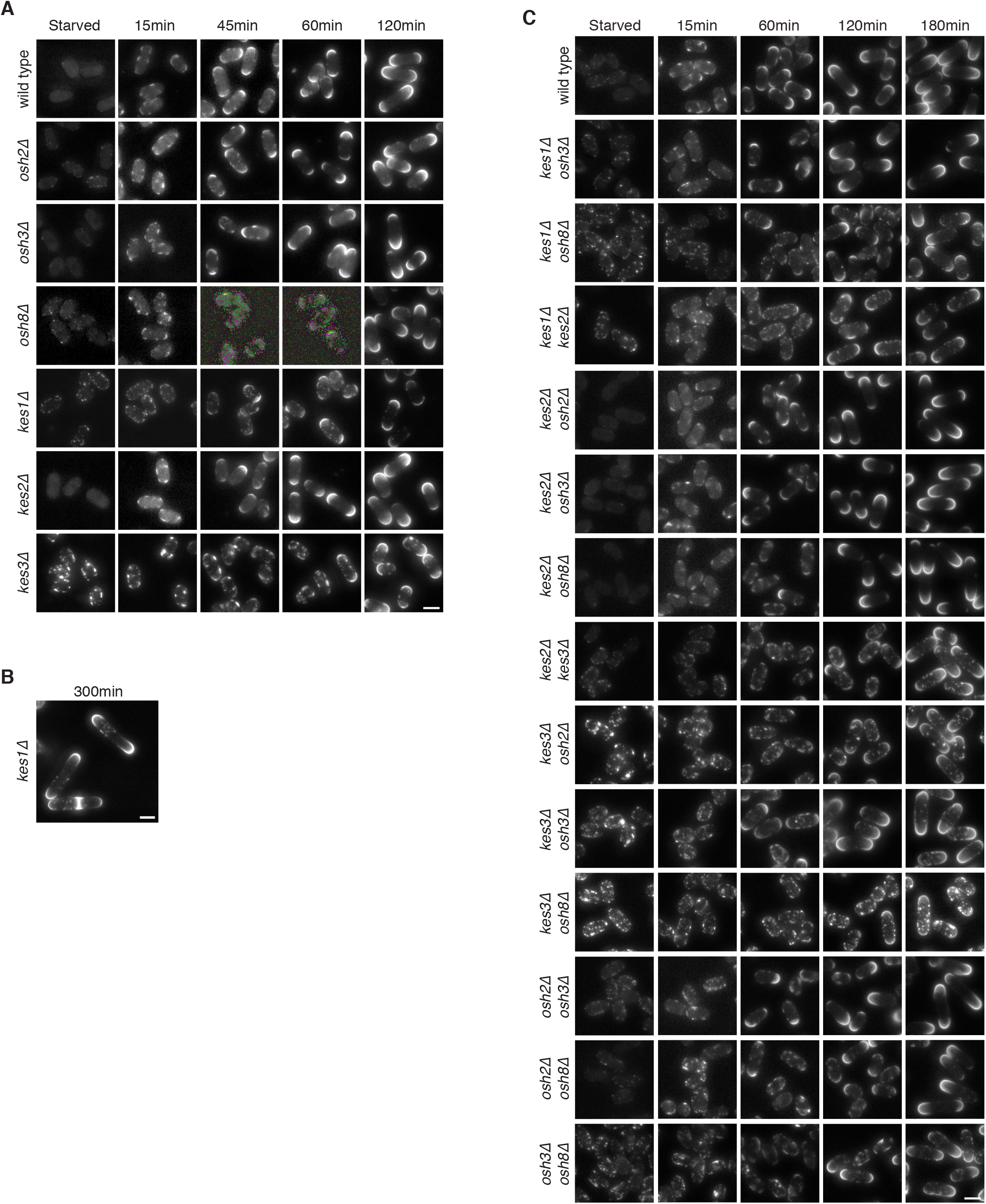
SE of cells carrying oxysterol-binding protein gene deletions. (**A**) SRM domain formation and polarisation during SE of wild type cells and of mutants stained with filipin and carrying gene deletions of oxysterol-binding proteins (*osh2*Δ, *osh3*Δ, *osh8*Δ, *kes1*Δ, *kes2*Δ and *kes3*Δ). (**B**) Asymmetric distribution of the residual filipin-stained sterol punctae in *kes1*Δ cells at the time of the first cell division during SE. (**C**) SRM domain formation and polarisation during SE of filipin-stained wild type cells and of all possible double mutant combinations shown in (A) and not shown in the main figures.

**Fig. S4:**
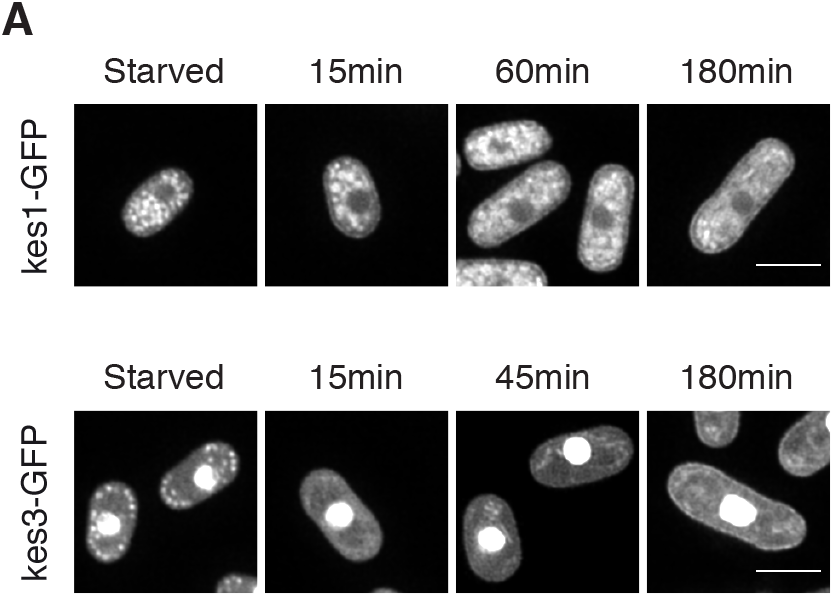
Non-polar localization of non-vesicular sterol delivery components. **(A)** The localization of GFP-tagged kes1p and GFP-tagged kes3p during SE.

**Fig. S5:**
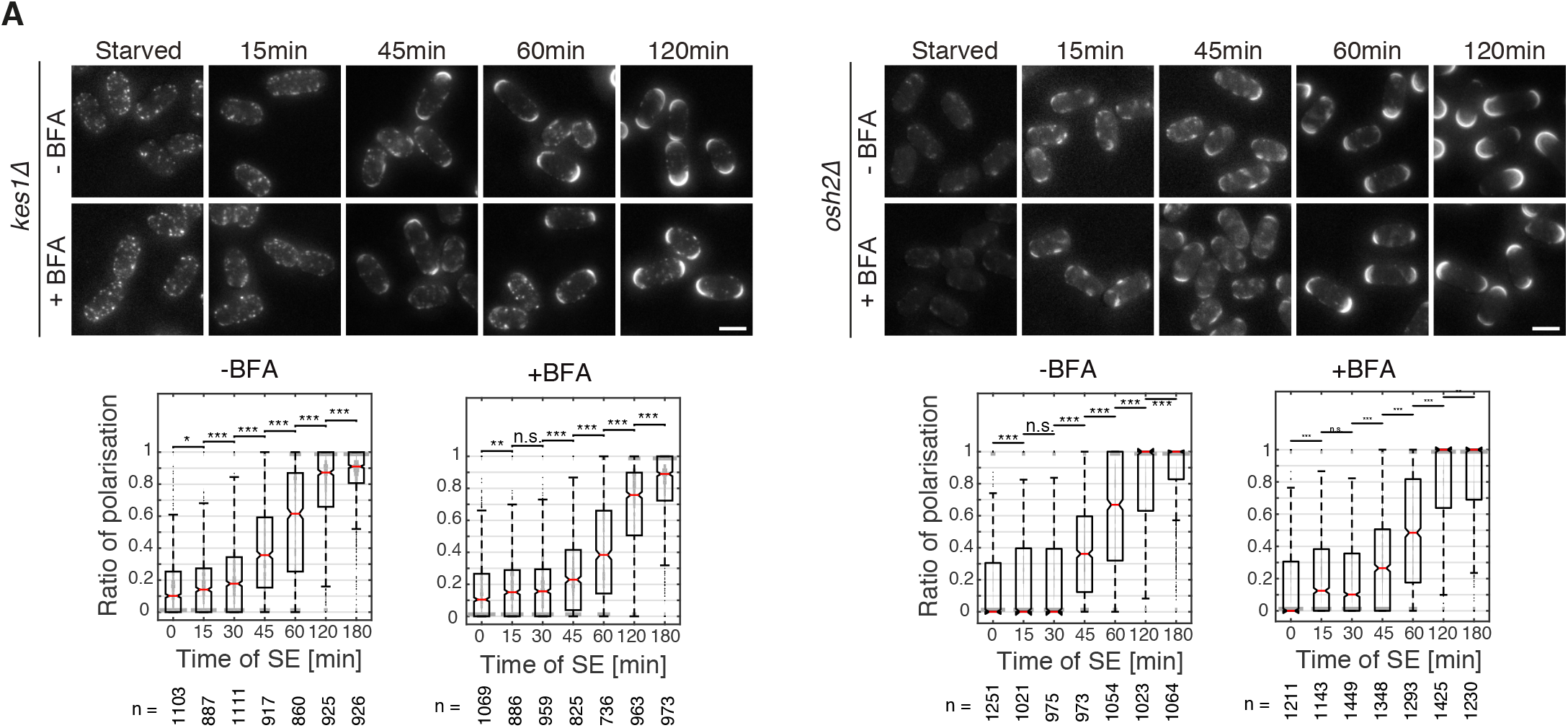
SE of *kes1*Δ and *osh2*Δ mutants with and without BFA. (**A**) SRM domain formation and polarisation during SE of filipin-stained *kes1*Δ and *kes3*Δ mutants treated with BFA dissolved in DMSO (+BFA) and DMSO alone (-BFA control). Notched box plots show the polarisation rates and cell lengths at different time points during SE with p-values. Scale bars: 5μm.

**Table S1:** List of strains used in this study. This table lists all strains used in the study. When labelled with “this study” strains were generated either *de novo* or by crossing existing strains, in which case the reference provides the origin of the parental strains.

| Strain | Genotype | Source | Adapted from |
| --- | --- | --- | --- |
| DB558 | *h-* | Leupold U. (1950) |  |
| DB681 | *h+ tea1Δ::ura4 ura4-D18* |  |  |
| DB1886 | *h- P3nmt1-GFP-tna1p::kanMX6* | Makushok et al. (2016) |  |
| DB4009 | *h- ypt2VN::ura4 ura4* | This study | Craighead et al. (1993) |
| DB4018 | *h- pob1-664* | This study | Toya et al. (1999) |
| DB4020 | *h+ cdc42-3::kanR* | This study | Tatebe et al. (2008) |
| DB4026 | *h- sec8-1* | This study | Wang et al. (2002) |
| DB4148 | *h- osh3Δ::kanMX6* | This study | Kim et al. (2010) |
| DB4168 | *h- sec3-916::HphR* | This study | Jourdain et al. (2012) |
| DB4178 | *h- osh2Δ::kanMX6* | This study | Kim et al. (2010) |
| DB4184 | *h+ kes1Δ::kanMX6* | This study | Kim et al. (2010) |
| DB4416 | *h+ kes3-HL-GFP::kanMX6* | This study |  |
| DB4826 | *h+ kes3Δ::kanMX6* | This study | Kim et al. (2010) |
| DB4847 | *h- kes1-HL-GFP::kanMX6* | This study |  |
| DB4848 | *h+ kes2Δ::kanMX6* | This study | Kim et al. (2010) |
| DB4850 | *h? pik1Δ::ura4 pREP41X-Ub-R-DHFRts-pik1* (referred as pik1-td cells) *leu1-32 ura4-D18* | This study | Park et al. (2009) |
| DB4854 | *h? kes1Δ::kanMX6 osh3Δ::kanMX6* | This study | Kim et al. (2010) |
| DB4857 | *h? kes1Δ::kanMX6 kes2Δ::kanMX6* | This study | Kim et al. (2010) |
| DB4869 | *h? pik1Δ::ura4 ade6-M210 leu1-32 ura4-D18, pREP41-2XeGFP-pik1* | This study | Park et al. (2009) |
| DB4879 | *h? kes2Δ::kanMX6 kes3Δ::kanMX6* | This study | Kim et al. (2010) |
| DB4885 | *h? sec3-916::HphR exo70Δ::kanMX6* | This study | Jourdain et al. (2012), Kim et al. (2010) |
| DB4939 | *h+ kes1Δ::kanMX6 kes3Δ::kanMX6* | This study | Kim et al. (2010) |
| DB4940 | *h? kes1Δ::kanMX6 cdc42-3::kanR* | This study | Kim et al. (2010), Tatebe et al. (2008) |
| DB4941 | *h? kes3Δ::kanMX6 cdc42-3::kanR* | This study | Kim et al. (2010), Tatebe et al. (2008) |
| DB4960 | *h? kes1Δ::kanMX6 tea1Δ::ura4 ura4-D18* | This study | Kim et al. (2010), Makushok et al. (2016) |
| DB5415 | *h? tea1HL-tagRFP::kanR* | This study | Makushok et al. (2016) |
| DB5416 | *h<sup>+</sup> leu2::sv40-GFP-atb2* | This study | Pardo M. & Nurse P. (2005) |
| DB5417 | *h<sup>+</sup> tea1HL-tagRFP::KanR cdc42-3::kanR* | This study | Makushok et al. (2016), Tatebe et al. (2008) |
| DB5418 | *h<sup>+</sup> leu2::sv40-GFP:atb2 cdc42-3::kanR* | This study | Pardo M. & Nurse P. (2005), Tatebe et al. (2008) |
| DB5781 | *h- osh8Δ::kanMX6* | This study | Kim et al. (2010) |
| DB5794 | *h- osh2Δ::kanMX6 osh3Δ::kanMX6* | This study | Kim et al. (2010) |
| DB5795 | *h- kes2Δ::kanMX6 osh3Δ::kanMX6* | This study | Kim et al. (2010) |
| DB5797 | *h- osh2Δ::kanMX6 osh8Δ::kanMX6* | This study | Kim et al. (2010) |
| DB5798 | *h- kes2Δ::kanMX6 osh2Δ::kanMX6* | This study | Kim et al. (2010) |
| DB5799 | *h- kes2Δ::kanMX6 osh8Δ::kanMX6* | This study | Kim et al. (2010) |
| DB5832 | *h- kes1Δ::kanMX6 osh8Δ::kanMX6* | This study | Kim et al. (2010) |
| DB5833 | *h- kes3Δ::kanMX6 osh8Δ::kanMX6* | This study | Kim et al. (2010) |
| DB5843 | *h- kes3Δ::kanMX6 osh2Δ::kanMX6* | This study | Kim et al. (2010) |
| DB5845 | *h- kes3Δ::kanMX6 osh3Δ::kanMX6* | This study | Kim et al. (2010) |
| DB5862 | *NAT::nmt81:kes1 NAT::nmt81:osh2* | This study |  |
| DB5864 | *h- osh3Δ::kanMX6 osh8Δ::kanMX6* | This study | Kim et al. (2010) |
| DB5943 | *h- kes1-HL-mGFP::kanMX6* | This Study |  |
| DB5944 | *h- kes3-HL-mGFP::kanMX6* | This Study |  |
| DB5949 | *h- osh2-HL-mGFP::kanMX6* | This Study |  |
| DB5973 | *h+ osh2Δ::NAT cdc42-3::kanR* | This study | Tatebe et al. (2008) |
| DB6008 | *h+ leu1::sec9-10 sec9Δ::NAT psy1-S1::ura4+ ura4-D18?* | This Study | YGRC/NBRP Japan |
| DB6026 | *h+ kes1Δ::kanMX6 leu1::sec9-10 sec9Δ::NAT psy1-S1::ura4+ ura4-D18?* | This Study |  |
| DB6027 | *h+ osh2Δ::kanMX6 leu1::sec9-10 sec9Δ::NAT psy1-S1::ura4+ ura4-D18?* | This Study |  |
| DB6048 | *h+ kanMX6::nmt81-kes3 leu1::sec9-10 sec9Δ::NAT psy1-S1::ura4+ ura4-D18?* | This Study |  |

**Table S2:**
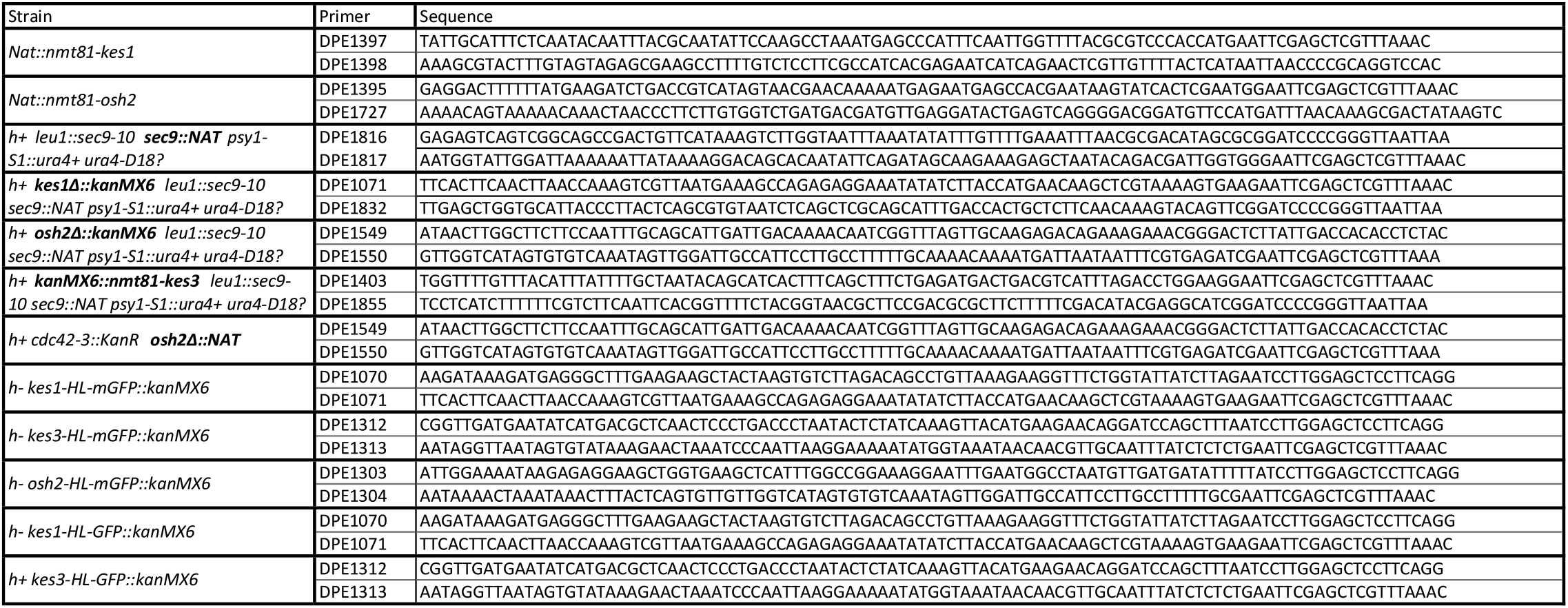
List of primers used in this study. This table lists the primer pairs used to generate the indicated strains. In strains with multiple insertions, the primers correspond to the insertion indicated in bold.

## Materials and Methods

### Yeast Culturing, Constructs and BFA treatment

For SE cells were cultured for 3 days at 25°C as described in Moreno et al. (1991) (*28*) in EMM with 0.5% glucose ^58^. Supplementing cells with EMM2 1:4 (starved culture:EMM2) triggered SE, which was done at 25°C unless stated differently. Starved temperature-sensitive mutants were preincubated at 36°C for 1h prior to SE at the restrictive temperature (36°C), which was triggered by adding pre-warmed EMM2. Cells were then kept at 36°C throughout SE. GFP-pikp, *pik1-td, nmt81:kes1 nmt81:osh2* and *nmt81:kes3 cdc42-3* strains were cultured in presence of thiamine during starvation and SE. The strains used are listed in Supplementary Information, Table 1.

Strains carrying ORP gene deletions were taken from the 5. edition Bioneer gene deletion library ^59^. Deletions were confirmed by colony PCR and auxotrophic markers were crossed out. Our ORP naming procedure is described in the Supplementary Information.

Gene deletions and protein-tagging were generated by homologous recombination as previously described ^60^. All primer pairs used for gene deletions or protein-tagging are listed in Supplementary Information, Table 2. *Nmt81::kes1* and *nmt81:osh2* were generated using the plasmid pFA6A-natMX6-P81nmt1. The strains *nmt81:kes1* and *nmt81:osh2* were then crossed to obtain the double shut-off strain. *Kes1*Δ and *osh2*Δ cells in the *psy1-S1 sec9-10* mutant background were generated using the plasmid pFA6A-kanMX6 and pFA6A-3HA-kanMX6 respectively. *Nmt81:kes3* in the *psy1-S1 sec9-10* mutant background was generated using the plasmid pFA6A-kanMX6-P81nmt1. *Osh2*Δ cells in the *cdc42-3* mutant background were generated using the plasmid pFA6A-3HA-Nat. Kes1p-GFP and kes3p-GFP strains were generated using plasmid pFA6A-HLGFP-kanMX6, which adds a previously published, 24aa HL linker (Happy Linker) in front of GFP(S65T) ^61^. Kes1p-mGFP, osh2p-mGFP and kes3p-mGFP strains were generated as previously described using a plasmid pFA6A-HL-GFP(A206K)-kanMX6, which contains a monomeric GFP ^41,56^.

BFA (LC Laboratories) was dissolved 50mg/ml in DMSO and used at 200μg/ml final concentration in the respective medium. Starved cells were pre-treated 1h with BFA. SE was initiated by adding BFA-containing EMM2 medium.

### Naming of ORPs

Pombase, the fission yeast genome database, lists six ORP protein family members (Fig. S2) ^34^. Two of them, kes1 and osh3, were previously given gene names. The other we named based on the annotated protein domains (Fig. S2) and overall protein homology to presumptive *S. cerevisiae* genes. Importantly, such overall protein sequence homology does not imply functional homology. This is most important in the case of kes3p, which based on sequence homology is listed in pombase as closest homolog of S. cerevisiae Osh7p. Osh7p was shown to bind phosphatidylserines in addition to sterols ^47^. Relevance as phosphatidylserine transporter was shown for the Osh7p homolog Osh6p. This Osh6p activity critically depends on defined features that are conserved in Osh7p and some other members of the ORP protein family. Fission yeast kes3p however, lacks all of these features showing that it is highly unlikely to bind phosphatidylserine. From this we conclude, that the closest functional homolog of kes3p in S. cerevisiae is the other ORP with high sequence homology, Kes1.

For our analysis we also considered *A. nidulans* orthologs as published by Bühler et al. 2015, as fission yeast *osh8* lacked a clear *S. cerevisiae* homolog and was most similar to *A. nidulans oshE* ^62^. Here, we simply applied successive numbering to fulfil *S. pombe* naming requirements.

### Microscopy/Image Analysis

Filipin (F9765, Sigma-Aldrich) staining is described in Takeda et al. (2004) and Makushok et. al. (2016) ^17,25^. Due to the use of higher quality DMSO and better storage vials, we could reduce the final concentration of filipin from 5μg/ml to 2μg/ml without compromising image quality. The cell wall was labelled with Rhodamine Grifonia Simplicifolia Lectin I (GFL I, BSL I), (Vector laboratories; 4μg/ml) as described in May and Mitchison (1986) ^63^. For imaging, samples of cells taken at defined time points of SE were placed on a Lectin-coated glass-bottom dish (Ibidi), stained and centrifuged at low speed. Imaging was performed on standard epifluorescence and spinning disc confocal microscopes as described in Makushok et al.^17^, using 40x (NA 1.3), and 100x (NA 1.4) oil objectives and CCD or sCMOS cameras for detection. Image stacks of 5 or 9 z-slices with 1μm or 0.5μm, respectively, steps were acquired for each acquisition channel. Image processing was done using ImageJ. For each z-slice of filipin and/or Rhodamine-lectin-stained cells we subtracted the background (rolling ball radius 100px) before performing a maximum projection. The image panels with filipin were processed qualitatively using optimized contrast settings for each panel individually, unless stated differently. There are two reasons for this: 1. More SRM domains are produced during SE, meaning that cells early in SE have a much lower filipin signal than cells later in SE. 2. The filipin signal intensity increases during the imaging process. This means that strains imaged later have a slightly higher filipin intensity than strains imaged earlier. However, all strains were imaged within 10min after adding filipin, at a time at which filipin is not yet damaging for the cells. For all other fluorescent images, routine image processing was done using comparable contrast settings.

Images in Fig. 6 and Fig. S4 were acquired as single plane images and deconvolved using Huygens deconvolution (Scientific Volume Imaging). P-values were calculated based on a Wilcoxon rank sum test ^64^.

### Single cell segmentation

To analyse and quantify SRM domain formation and polarisation dynamics, cells were stained with filipin and Rhodamine Grifonia Simplicifolia-Lectin I (Rhodamine-lectin). Image stacks of 5 z-slices with 1μm steps, or 9 z-slices with 0.5μm steps, were acquired for both fluorescence signals. In addition, a brightfield stack was also recorded. For each z-slice of filipin and Rhodamine-lectin-stained cells we subtracted the background (rolling ball radius 100px) before performing a maximum intensity projection. The Rhodamine-lectin signal was normalized using the *Enhance Contrast* plugin of ImageJ. For cell segmentation object probabilities were estimated using Ilastik software on the processed Rhodamine-lectin images ^65^. The generated cell probabilities were used in an adapted version of the previously described custom-made processing pipeline of CellProfiler ^66–68^.

Alternatively, the brightfield images were used for cell segmentation. A slice 1.5-2μm above and one the same distance below the focal plane were selected and the intensities of the top plane were subtracted from the bottom plane. The resulting image was corrected for background camera artefacts and used in a custom-made CellProfiler pipeline to detect cell outlines ^66^. Overlaying the boundaries of the segmented cells on images with the Rhodamine-Lectin- and or filipin staining was used to control segmentation quality. Artefacts were minor, showing that the segmentation procedure was very robust.

### Quantification of cell polarisation states

To quantify the polarisation state of individual *S. pombe* cells microscopy images were segmented as described above. For further analysis custom software was written in Matlab (MathWorks Inc.). A detailed description can be found in the supplementary material. In short: An estimate of the membrane was computed from the segmentation and local normals were fitted to the membrane curvature. Along these normals either mean or maximum intensities were measured. The corresponding intensity values for each pixel of the membrane curvature were interpolated to a reference curvature length and a designated signal processing and peak detection routine identifies regimes of high intensities guided by user-supplied parameters. These high intensity regions were classified as being at the poles or in between the poles of the fission yeast cells based on position information on the reference curvature. Each region was scored based on its size and mean intensity. A total polarisation score was then defined as the ratio of scores attributed to cell poles and to the cell regions in between the poles. The Matlab code is available on Bitbucket (https://bitbucket.org/Dreher/yeast-border-trace/).

### Measurement of cell eccentricity

The eccentricity is defined as the ratio of distance between foci and the major axis length of the inertia ellipse fit to the object. The value is between 0 and 1, where 0 corresponds to a circle-like and 1 to a line-like object. This was computed by Matlab’s built-in regionprops function (https://ch.mathworks.com/help/images/ref/regionprops.html).

