## Supplementary material for "Essential role of non-vesicular lipid transport in microtubule-controlled cell polarisation": Analysis_Software_Details

### <u>Automated analyses of SRM domain intensities and distribution</u>

We developed a Matlab routine to characterize fluorescent signal intensities on the cell membrane in relation to the circumferential position. The tool can be used for any fluorescent proteins or dyes and can be adapted to different cell sizes and, to some extent, to different cell shapes. This makes it a versatile analytical tool for use in various experimental approaches. The goal was to develop a simple tool that reliably measures the intensity distribution with reference to the fission yeast cell tip regions to provide a simple measure scoring phenotypes based on these distributions.

The underlying code is available on Bitbucket: https://bitbucket.org/Dreher/yeast-border-trace/. The published code is identical to the one we used for data analysis in our manuscript but has 2 additions: 1. It includes a graphical user interphase (GUI) making it very user friendly. 2. The GUI allows tweaking of the detection parameters. The data analysis in our manuscript used optimised, fixed parameters to guarantee a comparable analysis of the datasets.

In the following we describe the basis and rational behind the Matlab routine.

#### Algorithm overview

The core functionality was split into three main subroutines: 1) Measuring the signal intensity around the plasma membrane, 2) detecting the localisation patterns of signal intensities, i.e. tip regions versus side regions and small spots versus patches and 3) generating an output table. This was done to optimize re-computation in the workflow. The first step is experiment-independent and only needs two parameters describing properties of the segmentation pipeline used, and of its results. The second step is experiment-specific and requires tuning by the expert. With the division into these two steps, the intensity results do not need to be re-computed enabling fast result generation for different parameter combinations. The third step is optional. The output table enables the simultaneous analysis of multiple experiments since it can take different metadata from the filenames and generate according entries in an output table together with most of the results.

#### Measuring Border Intensities

The software expects the cell segmentation and the intensity or signal images in two separate folders, in which filepaths are given as input by the user. The images should be named in such a way that when sorted, corresponding images can be paired. This should be given by varying post- or prefixes included in the filenames (i.e. adding a unique number to the corresponding cell segmentation and signal images). Segmentation and intensity images are read and sorted by natural string sorting to assure matching images are taken, in case filenames do not include leading zeros for numerical information.

Components in contact with the image border are removed from the segmentation image to eliminate cropped cells. The image is then parsed to determine its type. We assume a binary image if there are only 0 and 1, or 0 and intmax values. Otherwise we assume a label image, in which each cell is a continuous area with a unique integer number. We want to differentiate the second case to match cell numbers with the defined label.

For binary segmentation images we compute the connected component structure typically used in Matlab imaging pipelines, by using bwconncomp. For label images, a custom helper function generates the same structure but with the correct order based on the labels. The image is converted to double without rescaling but a flag can be set to rescale the intensity range to a maximum of 1 (without rescaling the lower bound to 0). Then the membrane signal is computed for each cell with support for parallelization via Matlab's Parallel Computing Toolbox (MathWorks, "Parallel Computing Toolbox - MATLAB" Available at: https://ch.mathworks.com/de/products/parallel-computing. html).

Existing segmentation pipelines are typically geared towards a positive or negative contraction bias, i.e. the segmentation is always slightly larger or smaller than the actual membrane outline. To improve the capturing of the membrane ring that lies slightly inwards/outwards, the user can select a direction bias and an assumed membrane width M. For the inside bias the membrane consists of an M wide perimeter ring lying fully inside the segmentation and for the outside bias fully outside the segmentation accordingly. Both rings are symmetric with regard to the outer perimeter of the original segmentation. Since this can change the size of the assumed cell, the ring image is filled and a range of classical statistical properties are computed for the full cell. These are: centroid position, orientation, major and minor axis length, area, eccentricity, mean intensity of the object, mean intensity along the membrane, number of perimeter pixels, Euclidean length of perimeter. The membrane ring is then morphological thinned to a skeleton ring and an overlay image is saved for the user, both with the original and the shifted perimeter ring. The new boundary is traced as is the original perimeter, i.e. the binary image is converted into a sorted list of X-Y coordinates of the boundary. At this point three failure scenarios can arise, which are captured by logging and skipping the cell.

1. The resulting membrane image completely closes the cell without empty area inside. This will cause the skeletonisation to result in a single point or a very small area. In case the resulting skeletonisation is less than half of the original perimeter, it is assumed the cell's area collapsed.
2. In case the segmentation has a waist smaller than the assumed cell area, e.g. two cells in close contact have been segmented as one object, this results in the membrane ring self-intersecting and two boundary traces.
3. It could be that the original perimeter image does not result in a single boundary trace because of a badly conditioned segmentation.

Since the positional distribution of the membrane signal is a core interest, the boundary trace alone is not sufficient. A reproducible cell specific coordinate system is needed that orients the boundary with regard to the tip of the cells. We assume a clear orientation along the long axis of the cell. In case the cell has no long axis, i.e. is spherical, this will result in random orientations but in such a case even an expert would be unable to identify a tip without secondary information. For the long axis we take the orientation of the major axis of an inertia ellipse fit to the full cell image. From the centre of mass of the full cell, we project a line with said angle and detect the intersection points with the boundary. This gives us the two tip positions on the boundary. Since with the available information, the two tips are virtually impossible to distinguish from each other, we assume the one lying in the right half plane from the centroid to be the first one. This ensures reproducibility and makes it easier for the user to identify which part of the image was referenced when comparing images with numerical results. Minor coordinate points are also determined as intersection of a line emanating from the centre of the two major points perpendicular to the major axis with the perimeter.

The traced boundary vectors are then shifted so they start at the first major point, i.e. the centre of the right facing tip. The length of the boundary is computed as the Euclidean distance of line segments connecting the centre point of each boundary pixel. Then the membrane intensity for each boundary point is computed.

In theory it would be sufficient to simply take the signal image pixel intensities at each boundary pixel, but this is highly sensitive to the initial segmentation and small positional errors can lead to large variations. Instead the algorithm assumes that the identified boundary is close to the signal distribution of interest but not necessarily needs to match it. For each boundary pixel a small neighbourhood of the boundary is taken (typically 10 pixels) and a second order parametric curve is fitted to these N points. The curve fit is evaluated for N points tN = [0,1] for the parametric value and the one closest (in Euclidean distance) to the current pixel is determined. The normal vector at this point is computed by deriving the parametric curve and calculating the normal vector of the null space using the built-in null function.

An intensity profile is then measured along the normal vector starting from a point outward with the user defined membrane width and ending inward with this width. Both the mean and maximum value of this profile are then saved as intensity results for this perimeter point. The boundary fit and intensity profiling is then repeated for each pixel on the boundary. The results are stored in a structure together with the other statistical measures on the cells and saved in mat-File with the core user defined parameters to assist with reproducibility.

#### Detecting Intensity Patterns

The idea underlying this analysis tool is to identify membrane intensity patterns that are descriptive of different experimental conditions. After detecting and measuring the intensity profiles around the membrane of each cell, patterns must be identified. The most interesting parameters of these patterns are localization with regard to the cell shape, brightness, size and number/distribution for each cell. One experimental assumption is that the size of these patterns is not absolute, but relative to the membrane.

To make the results size relative, all 1D intensity profiles are interpolated to a fixed sample length. Optionally, the profiles can be filtered with a user defined filter to suppress noise and improve the detection routine. Implemented are options for Gaussian smoothing, moving-average, lowpass filtering, Savitzky-Golay filtering and discrete stationary wavelet transform. The helper function implements the correct circular padding for the data and parsing the necessary input parameters, while the filter implementations are available in Matlab.

For each re-sampled and smoothed intensity profile a peak detection routine is then run. The user can choose to correct for a baseline by shifting the profile such that the minimal y-value corresponds to 0. The peak detection routine is purposely modular. It offers a way to call any user definable peak detection routine. These can be supplied as Matlab function handle or function names in the search path. To this function the padded 1D intensity profile and two user parameter structures, FinderOptions and WidthOptions, are passed. The pipeline then expects four return values: peak values, peak locations, width (as matrix with left and right starting point) and the reference level for the width. Default is a bundled implementation based on the built-in findpeaks method but with a more powerful width measurement routine.

The determined peaks are classified as polar or side spots based on their width start and end point. If the width encompasses a tip point it is a polar spot, and if not, it is a side spot. For each spot a couple of characteristic measurements are saved to the output structure: the size or width of the spot, the mean and max intensity of the spot together with the full intensity vectors of each spot, the base value for width measurement and the position of the left width point, the peak and the right width. With all spots measured and classified the ratio of polarization and bi-polarity is computed for each cell.

#### Polarization Ratio and Bi-polarity Ratio

The polarization ratio should be a measurement of how much of the overall spot distribution is localized to the peaks. This definition alone is not sufficient for a mathematical definition, since there are a lot of free parameters. Simply basing the polarization ratio on the total intensity distribution that was localized to the poles relative to the rest turned out to be insufficient for certain experiments, where users wanted more control over how different characteristics, such as size or intensity, play into the polarization ratio. Depending on the imaging conditions, biological markers or underlying question, the importance of the observed criteria can shift, e.g. in some experiments the size distribution of the spots is weighted more importantly by the experimenter than the intensity values of the spots.

Hence for each spot, polar ($1...N_P$) and side ($1...N_S$) a score of importance is computed (S) and the ratio of polarization eq. (a) is then the ratio of scores of all polar spots relative to all spot scores. Each spot score is a multiplicative aggregation with individual weights, i.e. $S = X_1^{w1} X_2^{w2} \cdots X_n^{wn}$ .

This approach eliminates the burden of normalization from the user and allows for easy combination of values that are on different scales, e.g. intensities and size, while also being closer to the user assumption of importance of signal aggregates. Each spot is scored based on the formula eq. (b) which aggregates the mean intensity and the max intensity of the spot, as well as the size and smoothness (implemented as standard deviation of discrete differences of neighbours). By tuning the weights, the user can select the importance of these measures and by assigning a weight of zero completely remove their contribution from the scoring. Additionally, for each measure the user can decide whether the raw or smoothed intensity values should be used for the calculation, this is especially important for the max intensity and smoothness measure.

$$R_{pol} = \frac{\sum_{p=1}^{N_P} S_p}{\sum_{p=1}^{N_P} S_p + \sum_{s=1}^{N_S} S_s} \qquad \text{a}$$

with Sp as score of polar spots and SS as score of side spots, computed as:

$$S = \mu(I)^{w_1} \cdot \max(I)^{w_2} \cdot l^{w_3} \cdot \sigma(\Delta(I))^{w_4} \qquad \text{b}$$

$$R_{bi} = \frac{S_{p_1}}{S_{p_1} + S_{p_2}} \qquad \text{c}$$

Similarly, the bi-polarity ratio is defined as the ratio of one polar score relative to the sum of the two eq. (c). This allows the user to distinguish polarized cells with signal solemnly at one pole vs. equally distributed polar caps, corresponding to two different but important phenotypes. Since this score can only be defined for two detected polar spots, the bi-polarity ratio is capped for 1 polar spot at 0 and otherwise as NaN.

The results are again stored in a .mat file paired with the user given settings to allow for restarting computation from this step and preserving reproducibility.

**Output preparation**

At this point the main detection routine is finished and the data is available to the user as a collection of data structures, one for each cell with a selection of statistical measures, like size, major and minor axis length, eccentricity, circumference length and finally a structure holding the spot information, like number of polar and side spots, the polarity ratios defined above and the individual spot measurements, like size, left flank position, right flank position, intensity values, max intensity.

Since this tool was designed to be used also by people with limited scripting knowledge in Matlab, it includes a tool to create a user-friendly, table-like output that can be exported into other tools like Microsoft Excel or R. Besides reformatting the data into a table format, it provides a convenient way of transferring encoded information from the file name into a useful table column. It is a very common and useful practice in biology to encode additional information directly in the naming of the microscopy image files. For example, a typical file name might be: DD20161101_0_4184_nodrug_25_w1Reflector_1_11.tiff. Here the first part indicates a user and a date, something that is not strictly relevant to the experiment but the second bit the 0 indicates a time point in the experiment plan, the next number encodes a genotype used, the following string a treatment the specimen was subjected to and the following number a temperature. The information after that comes mostly from the imaging software and is trifling to the user. By passing a regular expression the user can translate this information to table columns (*Regular Expressions*, 1997. Available at: http://pubs.opengroup.org/onlinepubs/007908799/xbd/re.html). For the example file name this could look like this: DD\d{8}_(\d+)_(\d+)_(.*)_(\d\d )_.*.tiff. Each of the so-called groups in parenthesis are captured and translated into columns for all spots that were detected in that file. To generate meaningful table headers, a name for each group must be supplied in our case for example: 'Timepoint', 'Genotype', 'Treatment' and 'Temperature'. In case the user prefers, especially for string inputs, a numerical entry rather than the raw string, another input can be given with possible values, for which matches will be numbered in that order.

The function generates two output tables. One with a row entry for each detected spot, still containing all information of the cell it belongs to. The second table holds all the results for each cell, discarding individual spot information with a broader, more accessible overview, containing only aggregate information of the spots, like total number of polar and side spots and the two polarization ratios introduced above. Besides the decoding of the file name the only additional information this function computes is a score, labelled 'Death-Score', which can serve as an indication to exclude certain cells because they have likely died during the experiment. The idea behind the score is that healthy cells are expected to have only signal in the boundary region and their core region should be mostly indistinguishable from the background. The score is defined as the median of the z-score of the cells' intensity values in relation to the total image intensities. This summarizes how many standard deviations above the mean pixel intensities the median of all cell pixels lies, i.e. a cell should have only few pixels above the mean image intensity and hence a low death score, while a presumably dead cell with overall signal will have a high score. The z-score is used to make it independent from image acquisition settings and have an easily understandable dimensionless concept. This can help the user to deselect false information.

#### GUI Description

Our yeast border trace tool is a Matlab toolbox with full functionality via scripting in Matlab files. After initial testing it became clear that a user interface would provide a highly sought addition to the project, especially for the iterative tuning of the peak detection parameters.

Therefore, we designed a GUI with Matlab's GUIDE tool. It exposes all of the core functionality of the main spot finding step. The initial intensity detection relies on two parameters only that are defined by the segmentation routine and as such do not typically require tuning. The GUI does not expose the full workflow. It still requires adapting the main settings, like input files in a script, and call the computation pipeline with a parameter that one wants to utilize the GUI. It is only designed to speed-up the most interactive and difficult part, the tuning of peak detection routine and scoring weights. Once border intensities have been identified, the GUI will open up and expose the spot detection routine.

The main GUI windows is shown in Fig. 1. It displays a single cell result at a time. With the buttons on the bottom the user can move through image files and individual cells, while the current cell and image number are displayed. When the user quits the application, it will exit and return the parameters but not continue execution. With 'Batch execute' the current parameters will be used for one complete detection run, starting again from the first image, first cell.

On the top left half, the smoothed intensities are displayed together with markers indicating the identified polar and side spots and their respective peaks and width measurements. Since the width measurement of a Gaussian has to rely on a height value relative to peak or prominence, the width measurement bar corresponds to this height value, with local minima or other peaks restricting this measure indicated by vertical dashed lines.

In addition to the numerical measurement, the image of both the intensity signal as well as the segmentation are displayed to the right of the graphs. This allows to quickly identify important characteristics. For example, the user can check whether the peaks detected correspond with his visual spot classification or whether segmentation and signal image are matched correctly (added by displaying the file names above).

On the right half are the user interface elements associated with tuning the parameters. On the far right the scoring weights for the four different contributors to the two polarization ratios can be adjusted. On the bottom of the box the two ratios are displayed. They update when the user changes the weights above. With tick boxes the user can select whether the underlying intensity values should be the smoothed or unsmoothed. This can be important for the max intensity and smoothness since both are heavily affected by filtering of the curve.

The settings for both filtering and peak detection have been moved to their own small user interfaces reachable via their respective buttons. This has two main reasons: first the underlying computation is too slow for a live and interactive user interface with reasonable response time and secondly both of these are purposely designed in a modular fashion. That way an advanced user can expand the software with additional filters or a different peak detection without the need for extensive modification. This approach was kept for the GUI such that an expansion can be provided together with a custom UI without having to touch the main functionality and code.

The filter GUI as shown in Fig. 2, consists of a simple drop-down menu with all supported filters and a changing parameter interface below. This adapts according to the selected filter from simple one-, to two-parameter fields, to an entry table with up to 8 parameters for wavelet filtering. The displayed dialogue is modal, i.e. the user can only interact with the filter GUI while it is open.

In the same way the peak detection GUI in Fig. 3 consists of simple entry fields for all parameters combined with drop-down menu displaying categorical settings. Parameters are grouped into the peak finding routine using findpeaks, exposing all meaningful parameters and width measurement for quantifying the peak width.

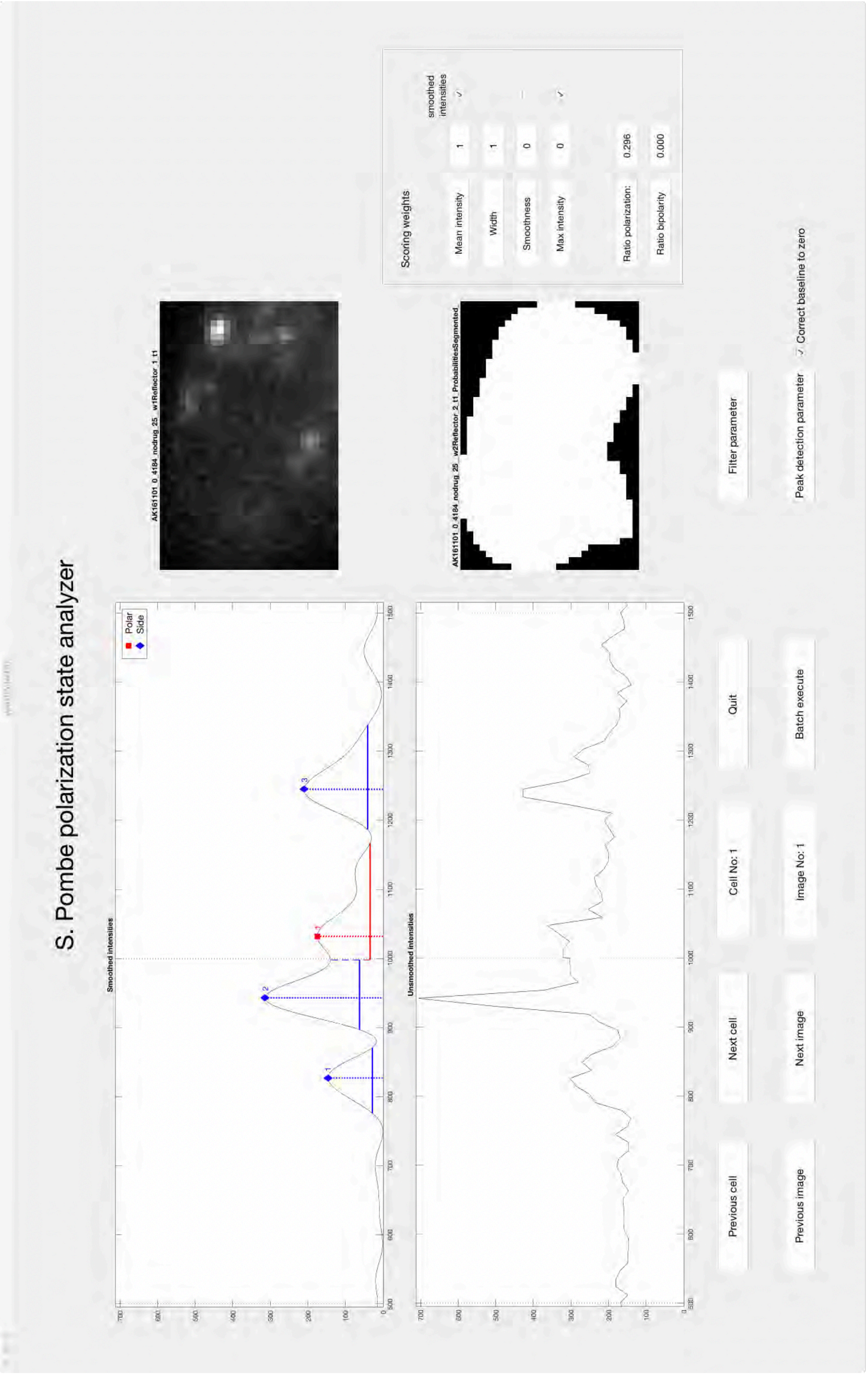


**Figure 1:** Main User Interface Yeast Border Trace Tool

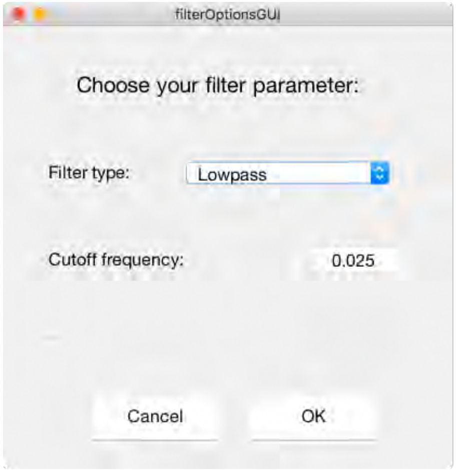


**Figure 2:** Filter Interface

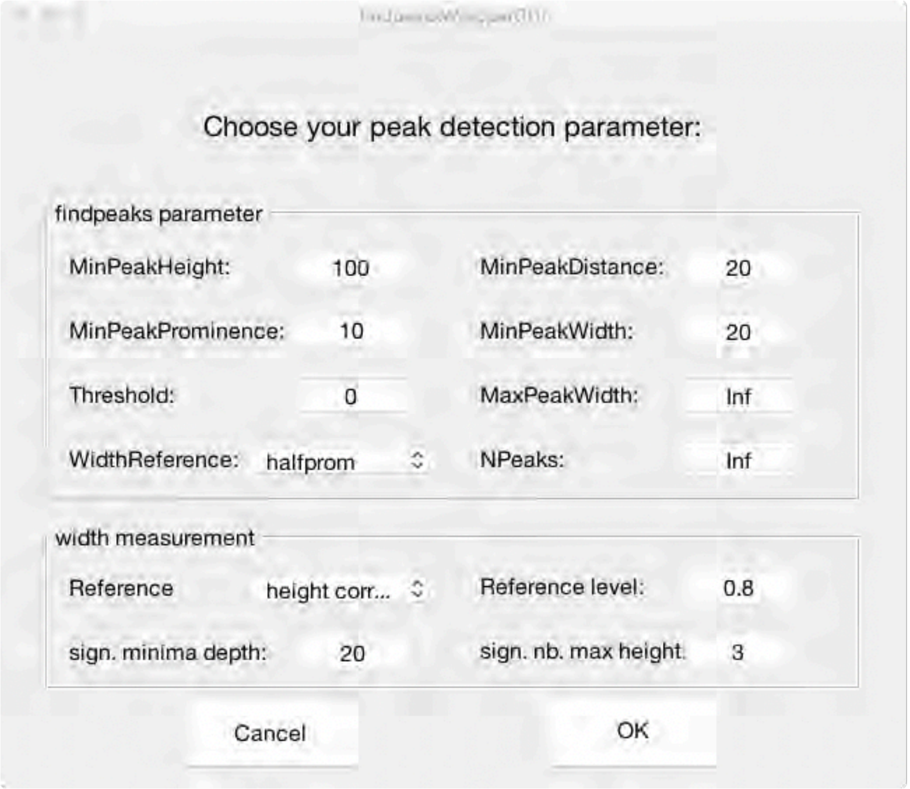


**Figure 3:** Peak Detection Interface
